# Effective control of a viral disease with a high transmission rate through selective predation

**DOI:** 10.1101/2020.09.24.311563

**Authors:** Yonggui Chen, Muhua Wang, Zhong Zhao, Shaoping Weng, Jinchuan Yang, Shangyun Liu, Chang Liu, Fenghua Yuan, Bin Ai, Haiqing Zhang, Mingyan Zhang, Lirong Lu, Kai Yuan, Zhaolong Yu, Bibo Mo, Zhiwei Zhong, Luwei Zheng, Guocan Feng, Shengwen Calvin Li, Jianguo He

**Affiliations:** State Key Laboratory for Biocontrol, School of Marine Sciences, Sun Yat-sen University, Guangzhou 510275, China; Southern Marine Science and Engineering Guangdong Laboratory (Zhuhai), Zhuhai 519000, China; School of Mathematics and Computational Sciences, Sun Yat-sen University, Guangzhou 510275, China; School of Life Sciences, Sun Yat-sen University, Guangzhou 510275, China; Maoming Branch, Guangdong Laboratory for Lingnan Modern Agricultural Science and Technology, Maoming 525435, China; University of California-Irvine School of Medicine, Children’s Hospital of Orange County, Orange, CA 92868-3874, USA

## Abstract

Due to the limited understanding of the characteristics of predator-pathogen-prey interactions, few attempts to use selective predation for controlling diseases in prey populations have been successful. The global pandemic of white spot syndrome (WSS), caused by white spot syndrome virus (WSSV), causes devastating economic losses in farmed shrimp production. Currently, there is no effective control for WSS. Here, we determined the transmission dynamics of WSSV and the feeding ability and selectivity of fish on healthy, infected and dead shrimp by experiments and mathematical modeling. Accordingly, we developed a novel and convenient shrimp cultural ecosystem, which that effectively prevented WSS outbreaks, by introducing aquaculture fish species. This provides a scheme for developing control strategies for viral diseases with high transmission rate.

---

In nature, predators prefer to eat infected prey, and it is predicted that selective predation on infected individuals can reduce the prevalence of diseases in the prey population (*1, 2*). The effectiveness of selective predation in achieving disease prevention is determined by the interplay of several factors (*3-5*). Without a comprehensive understanding of the characteristics of predators, pathogens and prey, there have been few successful attempts to employ selective predation to control diseases in prey populations, especially diseases with high transmission rates (*6-8*).

Aquaculture is one of the major sources of animal protein, and shrimp are the primary global aquaculture species (*9*). However, white spot syndrome (WSS), which is caused by white spot syndrome virus (WSSV), has led to considerable economic losses to the global shrimp aquaculture industry (*10*). At present, there are no effective prevention and control strategies for WSS, although several methods have been tested in farmed shrimp systems (*11-14*). WSS pandemics primarily occur with the sequential transmission of WSSV from healthy shrimps consuming WSSV-infected shrimps to other healthy shrimps (*15, 16*), lacking a predator to remove the infected individuals from the high density single-species culture system (**Fig. S1**). The prevalence of diseases in the prey population is positively correlated with the disease transmission rate but is negatively correlated with the predation pressure and predator selectivity (*17, 18*). Thus, to develop a shrimp culture system that controls WSSV outbreaks through selective predation, we studied WSSV transmission dynamics in a Pacific white shrimp (*Litopenaeus vanmamei*) population and elucidated the feeding ability and selectivity of various species of aquaculture fishes on shrimp.

To determine the transmission dynamics of WSS, we performed studies on the relationships among the body weight of one initial WSSV-infected shrimp, number of deaths, and death time distribution. One piece of WSSV-infected dead shrimp infected several healthy shrimps with the same bodyweight via ingestion. The infected number of shrimps in the groups exhibiting average body weights of 1.98 g, 6.13 g, and 7.95 g were 57.3, 64.7 and 71.3, respectively (**Fig S2 and Table S1**). This suggests that the transmission rate of WSSV is extremely high, and the basic reproduction number (*R_0_*) of WSSV increases with the body weight of WSSV-infected dead shrimp. Time to death was consistent across the three groups of body weights for WSSV-infected shrimp, with the majority of death occurring on the third to sixth day and the peak number of deaths occurring on the fourth and fifth days (**Figs. 1A and S3, and Tables S2 to S4**). A mathematical model (Model 1) was developed to describe the transmission dynamics of WSS (**Figs. 1A and 1B, Fig. S1, Supplementary Text**). In addition, the changes in live and dead shrimp numbers during WSSV transmission were determined by artificial infection experiments. The number of live shrimps began to decrease after 2 days of WSSV infection and drastically decreased after 4 days of WSSV infection (**Fig. 1C**). The dynamic changes of healthy, infected, and dead shrimp could be expressed by a mathematical model (Model 2) (**Figs. S1, S4 and S5, Supplementary Text**). Model 2 predicts that it is possible to cut off the transmission route of WSSV by removing infected and dead shrimp, but the time window of prevention is approximately 2 days. Thus, the fish utilized as predators must ingest infected and dead shrimp promptly and continuously.

**Fig. 1.**
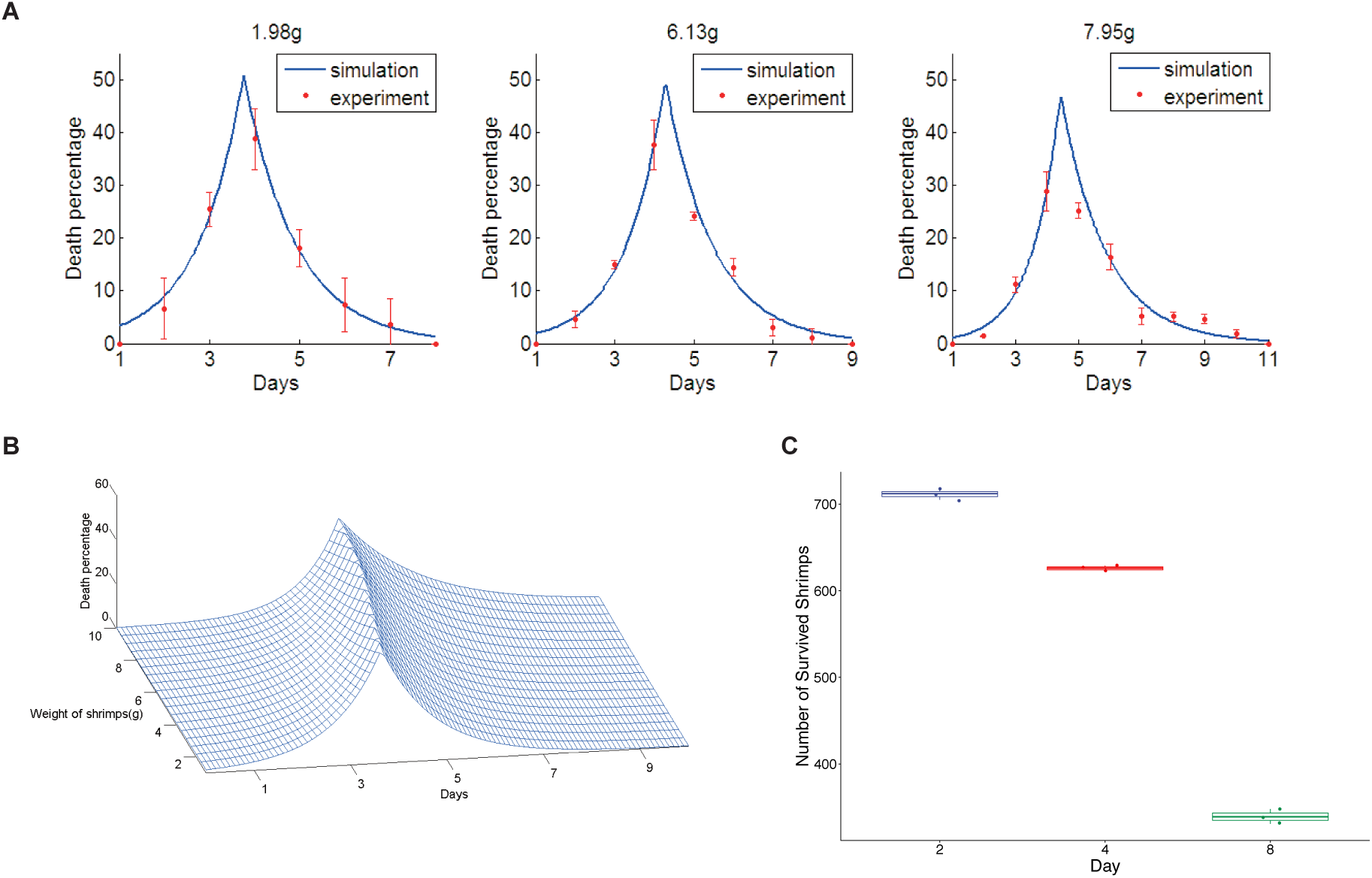
Transmission dynamics of WSSV. **(A)** Daily death percentage of shrimp populations with initial WSSV-infected shrimp of different body weights. The red points with error bars are the results from experiments, and the solid blue lines are the results of model 1. **(B)** The illustration shows the death percentages of infected shrimp with different body weights each day. We estimate the parameters in model l and draw the above 3D surface to show the relationship between the death percentage concerning shrimp weights and time. For all weights of shrimp, the death percentage rises at the beginning and then drops. The peak time for death was approximately the fourth day. **(C)** The relationship between the number of surviving shrimp and days after healthy shrimp were cocultured with WSSV-infected shrimp. Means and standard errors are shown.

To identify the fish species for controlling WSSV transmission, we examined the feeding ability and selectivity of diverse aquaculture fish species that ingest Pacific white shrimp, including grass carp (*Ctenopharyngodon idella*), African sharptooth catfish (*Clarias gariepinus*) (hereafter referred to as catfish), red drum (*Sciaenops ocellatus*), and brown-marbled grouper (*Epinephelus fuscoguttatus*). The brown-marbled grouper did not ingest dead shrimp continuously, and its daily dead shrimp ingestion rate was low (**Table S5**). The daily dead shrimp ingestion rates of grass carp, catfish and red drum were 8.26%, 4.99%, and 11.63%, respectively (**Tables S6 to S8**). The daily healthy shrimp ingestion rates of grass carp, catfish and red drum were 2.09%, 1.01% and 6.04%, respectively, indicating the relatively high healthy shrimp ingestion rate of red drum (**Tables S9 to S11**). Grass carp and catfish have high feeding selectivity of dead shrimp over infected and healthy shrimp (**Figs. 2A and S6, and Tables S12 and S13**). Additionally, grass carp and catfish move quickly and swallow infected and dead shrimp completely. These characteristics suggest that grass carp and catfish have high feeding selectivity and ability, which can cut off the WSSV transmission route in which healthy shrimp ingest infected dead shrimp.

**Fig. 2.**
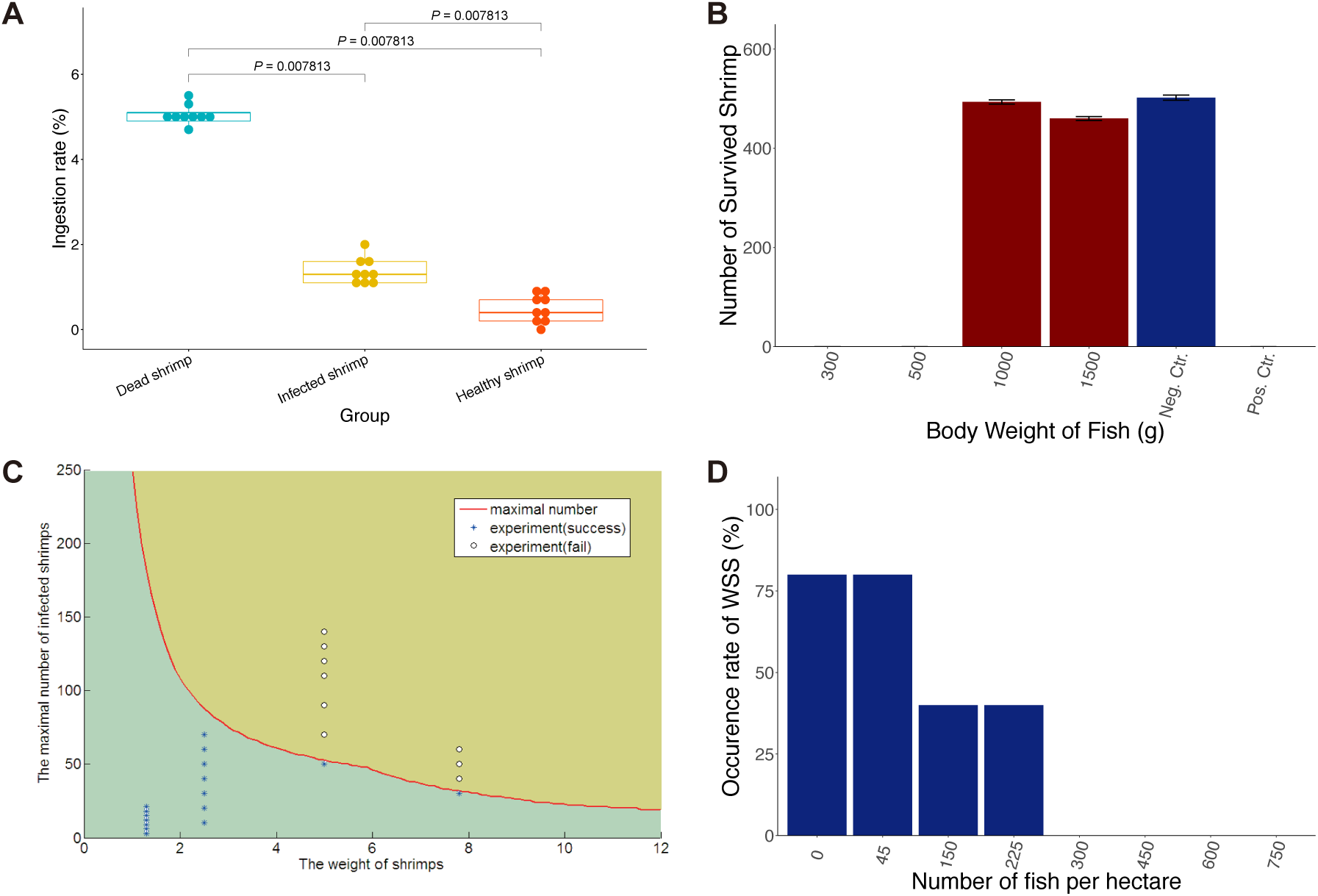
Specification of grass carp for control of WSSV transmission. **(A)** Feeding selectivity of grass carp on dead, infected (endopod and exopod removed), and healthy shrimp. The diseased shrimp infected with WSSV had reduced activity and died within one day. The activity of shrimp was reduced after the endopod and exopod were removed. Thus, shrimp with endopod and exopod removal were utilized to resemble WSSV-infected shrimp. *P*-values (permutation test, paired) were labeled. Grass carp ingested significantly more dead shrimp than infected (endopod and exopod removed) and healthy shrimp. **(B)** The effect of the different body weights of grass carp on the control of WSS outbreaks. Means and standard errors are shown. **(C)** Capacity of 1-kg grass carp to control WSS. Blue asterisks represent the number of infected shrimps successfully controlled by one 1-kg grass carp, while clear circles represent the number of infected shrimps that failed to be controlled by one 1-kg grass carp. The red line is the simulated highest value of one 1-kg fish that can control the number of infected shrimps with different body weights based on model 3. **(D)** The relationship of the number of cocultured grass carp and the occurrence rate of WSS. More than 300 grass carps of approximately 1.0 kg per hectare can completely control the outbreaks of WSS, but fewer than 225 grass carp cannot fully control outbreaks of WSS.

The capacity and specification of fish were determined for effective control of WSS. First, we identified the minimum body weight of cocultured grass carp and catfish. Four experimental ponds were set up in which 600 healthy and 3 WSSV-infected shrimp with body weights of 5 g were cultured with one grass carp of different body weights. After 13 days in culture, the ponds cocultured with one grass carp weighing 0.3 kg, 0.5 kg, 1 kg, and 1.5 kg showed shrimp survival rates of 0, 0, 82.4%, and 79.4%, respectively (**Fig. 2B and Table S14**). This finding indicates that the minimum body weight of grass carp for effective control of WSS is 1 kg. Furthermore, experiments showed that coculturing catfish with body weights greater than 0.5 kg could effectively control WSS transmission in the shrimp population (**Fig. S7 and Table S15**). Second, we determined the capacity of grass carp for WSS prevention in shrimp populations of different shrimp body weights based on experimental results. In a 10-m^2^ pond with 750 shrimps, one 1-kg grass carp could control 70 pieces of 2.5 g WSSV-infected shrimp, 50 pieces of 5.0 g WSSV-infected shrimp, or 30 pieces of 7.8 g WSSV-infected shrimp, which is consistent with the result of the mathematical simulation (Model 3) (**Fig. 2C, Fig. S1, Table S16, Supplementary Text**). This result suggests that the capacity of grass carp to control WSS is negatively correlated with the body weight of shrimp. Thus, releasing fishes in the early stages of shrimp production may improve their capacity to control WSS. Finally, the minimum stocking quantity of grass carp to effectively control WSS was determined. We released 45, 150, 225, 300, 450, 600 and 750 grass carps/hm^2^ of bodyweight of 1 kg∼1.25 kg into the 40 ponds (13.57 hm^2^) at a demonstration farm (**Figs. 2D and S8, and Table S17**). When grass carp stocking quantities were greater than or equal to 300 grass carp/hm^2^, the success rate of WSS control was 100%.

Next, we translated the knowledge obtained from these experiments to an applied technology scale. First, the effectiveness of controlling WSSV transmission by coculturing shrimp with grass carp was tested in two experimental zones at a farm in Maoming, Guangdong Province, China (Farm 1) in 2011 (**Fig. 3A**). In the 18 ponds (6.03 hm^2^) of zone A, we cultured 9×10^5^/hm^2^ of shrimp for 20 days and then introduced 317–450/hm^2^ of grass carp with average body weights of 1 kg. We did not observe WSS outbreaks in 17 ponds and harvested 7,332 ± 2,059 kg/hm^2^ in 110 days of culture (**Fig. 3B and Table S18**). One pond was unsuccessful due to pathogenic bacterium (Vibrio) infection. Shrimps were cultivated without grass carp in 28 ponds (11.30 hm^2^) in zone B. A total of 20 ponds in zone B had WSS outbreaks, resulting in an average yield of 1,844 ± 2,304 kg/hm^2^. In 2012, we switched zones A and B, cultivating shrimp with grass carp in zone B but without fish in zone A (**Fig. 3B and Table S18**). In all ponds of zone B, we did not observe WSS outbreaks and harvested 8,587 ± 1,655 kg/hm^2^ in 110 days of culture. However, the average yield of zone A was 1,953 ± 2,188 kg/hm^2^ due to WSS outbreaks, which occurred in 12 of 18 ponds. Second, to evaluate the effectiveness of controlling WSSV transmission by coculturing shrimp with catfish, two experimental zones were designed at a farm in Qinzhou, Guangxi Province, China (Farm 2) in 2011 (**Fig. 3C**). In zone A, we cultivated 7.5×10^5^/hm^2^ of shrimp in 38 ponds (21.20 hm^2^) for 10 days and then introduced 525–750/hm^2^ of catfish with average body weights of 0.5 kg. Shrimp were cultivated in 57 ponds (67.00 hm^2^) without fish in zone B. In zone A, we did not observe WSS outbreaks in any of the ponds and harvested 8730 ± 1187 kg/hm^2^ in 110 days of culture (**Fig. 3E and Table S19**). However, WSS outbreaks occurred in 53 ponds of zone B, resulting in an average yield of 450 ± 1420 kg/hm^2^. In 2012, we split zone B into zones B1 and B2. Shrimp were cultivated with catfish in 38 ponds of zone A and 25 ponds (27.00 hm^2^) of zone B1, while shrimp were cultivated without fish in 32 ponds (40.00 hm^2^) of zone B2 (**Fig. 3D**). We did not observe WSS outbreaks and harvested 9628 ± 1471 kg/hm^2^ in zone A and 6375 ± 1000 kg/hm^2^ in zone B1. However, WSS outbreaks were observed in 29 of 32 ponds in zone B2, resulting in an average yield of 500 ± 1900 kg/hm^2^ (**Fig. 3E and Table S19**). Moreover, releasing grass carp and/or catfish effectively controlled the WSS outbreak and substantially increased shrimp production at Farm 1 between 2013 and 2019 (**Fig. 3F and Data S1**).

**Fig. 3.**
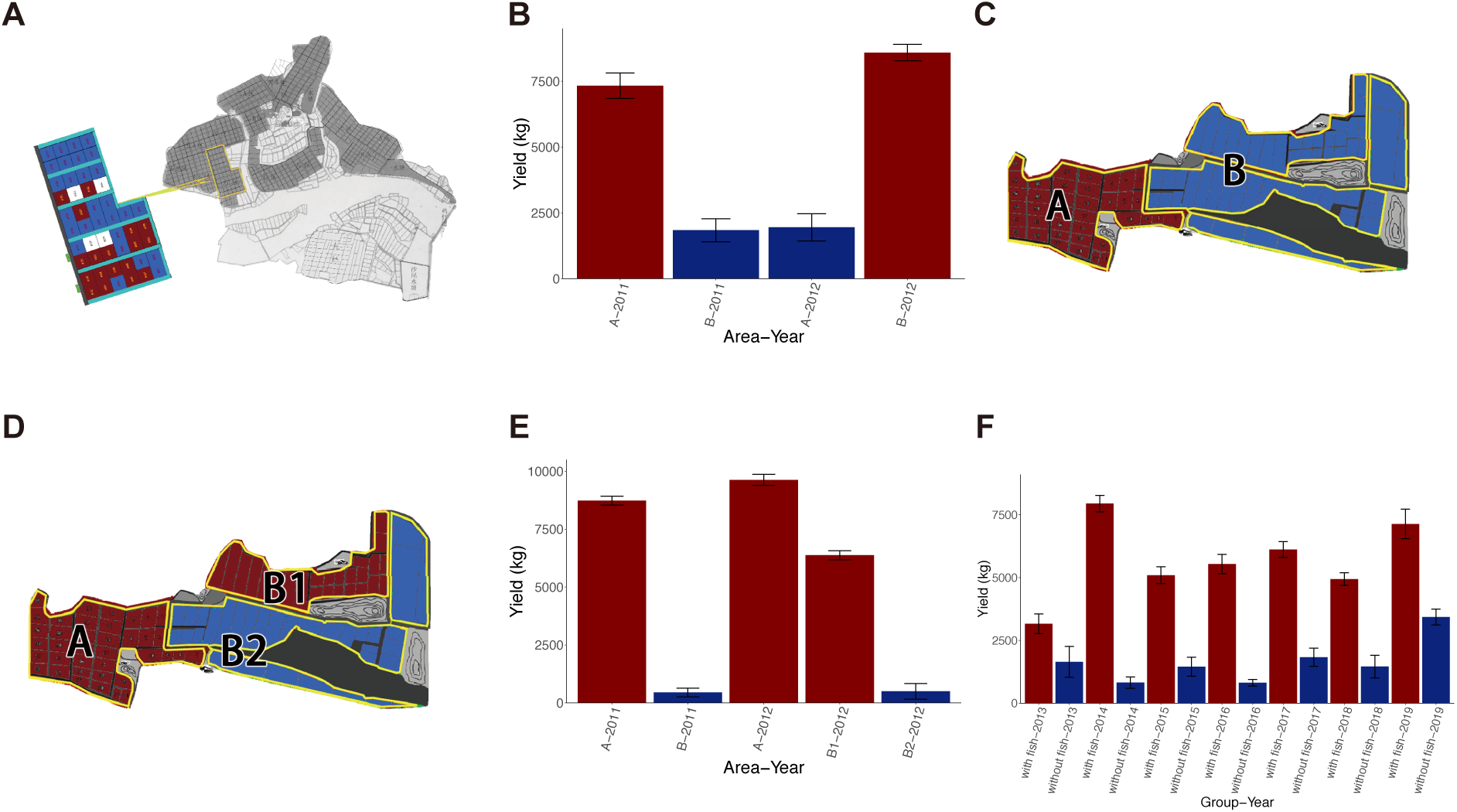
Control of WSS in shrimp production by fish. **(A)** The design of the field study for the control of WSS using grass carp. The satellite map of the farm at Maoming, Guangdong Province, China (Farm 1) is shown. The 46 experimental ponds were divided into zone A (red) and zone B (blue). In 2011, shrimp were cultured with grass carp in ponds in area A, while shrimp were cultured without fish in ponds in area B. In 2012, shrimp were cocultured with grass carp in area B but without fish in area A. **(B)** Total yield of shrimp production in ponds with (red) or without grass carp (blue) at Farm 1. Means and standard errors are shown. **(C)** The design of the field study for the control of WSS using catfish in 2011. The satellite map of the farm in Qinzhou, Guangxi Province, China (Farm 2) is shown. The 95 experimental ponds were divided into zone A (red) and zone B (blue). Shrimps were cultured with catfishes in the ponds in area A, while shrimps were cultured without fishes in the ponds in area B. **(D)** The design of the field study for the control of WSS using catfish in 2012. Shrimps continued to be cultured with catfishes in the ponds in area A, while area B was divided into two groups: shrimps were cultured with catfishes in the ponds in area B1, and shrimps were cultured without fishes in the ponds in area B2. **(E)** Total yield of shrimp production in ponds with (red) or without catfish (blue) at Farm 2. Means and standard errors are shown. (F) Total yield of shrimp production in ponds with (red) or without fishes (blue) at Farm 1 from 2013 to 2019. Means and standard errors are shown.

By elucidating the transmission dynamics of WSSV, we developed a cultivation system to control disease outbreaks in shrimp populations by restoring the interactions of predators and prey. In addition, the species, body weight, and density of cocultured fishes were determined by determining the feeding ability and selectivity of fishes on shrimp through experiments. To the best of our knowledge, this report describes the first artificial system that successfully controlled viral disease with a high transmission rate through selective predation. Furthermore, the use of aquaculture fish as predators results in the production of shrimp and fish simultaneously in the system. The results of this study highlight the significance and provide a scheme of elucidating the characteristics of predators, pathogens, and prey to develop disease control strategies through selective predation.

## Funding

This project was fund by the National Natural Science Foundation of China (No. U1131002), the Chinese Agriculture Research System (No. CARS-47), the National Key Technology R&D Program (2012BAD17B03), the Special Fund for Agro-scientific Research in the Public Interest (No. 201103034), and the National Basic Research Program of China (No. 2012CB114401).

## Author contributions

J.G.H. conceived of the project and designed research, S.C.L. and G.C.F. helped the proceeding. Y.G.C. and S.P.W. were the lead coordinators for laboratory and field study; Y.G.C, J.C.Y., S.Y.L., C.L., F.H.Y., Z.W.Z., H.Q.Z., M.Y.Z., L.R.L, and K.Y. performed the experiments; Z.L.Y., B.B.M. and L.W.Z performed the field study; M.W, and B.A. analyzed the data; Z.Z. and G.C.F. performed the mathematical modeling; M.W, S.C.L. and J.G.H. wrote the manuscript with the contribution from all authors.

## Competing interests

Authors declare no competing interests.

## Data and materials availability

All data is available in the main text or the supplementary materials.

## Notes

### Competing Interest Statement

The authors have declared no competing interest.

### Summary of Updates

The title of the manuscript has been revised. The writing of the manuscript has been improved for clarity and fluency.

