## Supplementary Materials for "Effective control of a viral disease with a high transmission rate through selective predation"

**Materials and Methods**

The relationship among the body weight of one initial WSSV-infected shrimp, number of deaths, and death time distribution.

To study the WSSV transmission rate in cultured shrimp populations through healthy shrimp ingestion of WSSV-infected dead shrimps, three groups of body weight 1.98 ± 0.03g, 6.13 ± 0.16g, and 7.95 ± 0.13g, respectively, were used. Each group consisted of three replicates and one control. Each group consisted of 430 shrimps, in which 30 shrimps were randomly selected for screening performed by a two-step WSSV PCR assay, showing negative. The remaining 400 shrimps were divided equally to four aquariums, and each aquarium consisted of 100 shrimps (Three replicates and one control). All 12 shrimp aquariums were designed with the same dimension of 220 cm × 60 cm × 80 cm, water volume of 0.5 $m^3$, temperature of 25 ~ 28 °C, and salinity of 8. One hundred shrimps were placed into each aquarium and quarantined for seven days to ensure no dead shrimps. In each group (four aquariums) of the similar body weight, three replicate aquariums were separately put in with one piece of artificially WSSV-infected dead shrimp of the same body weight, and the fourth aquarium was put in with one piece of frozen dead shrimp (WSSV-free) of the same body weight as a control. Shrimps were fed once a day with artificial feed (ratio of 2% body weight). Shrimp feces were timely removed, and 50% of the water was exchanged with water (salinity of 8) every day. To prevent repeated infections of shrimp, from the second day of infection, shrimps were observed every 10 minutes to screen and remove dying shrimps. When moribund (dying) shrimps appeared, they were promptly removed to prevent healthy shrimps from infection by eating these dying shrimps but not the initial infected dead shrimp. These criteria determined moribund (dying) shrimps: capable of pleopod activity, but no response (no mobility) by glass rod agitation. This screen was continued until dying shrimps no longer appeared three days after the appearance of the last dying shrimp (~ 15 days). Dying and remaining live (survived) shrimps were subjected to the one-step WSSV PCR assay, showing WSSV-positive in moribund shrimps, and negative in live shrimps. Based on the above experimental results, a mathematical model of the relationship among time distribution, number of deaths and body weight of WSSV infected shrimp was established (Model 1).

The dynamic changes of live, infected, and dead shrimps during WSSV transmission

To determine the changes of numbers of live and dead shrimps during WSSV transmission, nine cement ponds (size of 10 $m^2$, water volume of 5 $m^3$) were set up.

Regarding the density of 7.5x$10^5$-shrimps/$hm^2$ in shrimp farming production, 750 WSSV-free shrimps (average body weight of 7.9 g) were delivered into each concrete pond. Shrimps were quarantined for seven days and then subjected to the feeding of 30 artificially WSSV-infected shrimps in each concrete pond. Shrimps were fed with artificial feed every day (ratio of 2% body weight). The number of survived shrimps was determined in three of the concrete ponds on the 4th, 6th, 8th day post-infection, respectively. Dead shrimps were assayed by using the one-step PCR, showing WSSV-positive. Based on the model 1, we established a mathematical model (Model 2) to describe the dynamic changes of healthy, infected, and dead shrimps during WSSV transmission.

The ratio of dead shrimp ingestion of fishes

To determine the rate of dead shrimp ingestion of brown-marbled grouper (*Epinephelus fuscoguttatus*). Three cement ponds were set up with the same dimensions of 315cm × 315cm × 120cm, water volume of 5 $m^3$, and salinity of 20. Three brown-marbled groupers with similar body weight were put in each pond. The average body weights of groupers put in each of the three ponds were about 0.03 kg, 0.15 kg, and 0.30 kg, respectively. And the total body weights of fishes in each of the three ponds are 0.098 kg, 0.501 kg, and 0.999 kg, respectively. The fishes were raised for four days before the experiment started. During the five days of experiment, fishes were fed with dead shrimps. The mean weight of dead shrimps used in the experiment is 5.9 g. Each day, the total body weight of dead shrimps that were ingested by fishes in each pond was calculated by subtracting the total body weight of dead shrimps remained in the pond from the total weight of dead shrimps put in the pond. The shrimp ingestion rate of fish is quantified by the daily ingestion rate (total body weight of ingested shrimps per day / total body weight of fishes).

To determine the rate of dead shrimp ingestion of grass carp (*Ctenopharyngodon idellus*). Three cement ponds were set up with the same dimensions of 315cm × 315cm × 120cm, water volume of 5 $m^3$, and salinity of 5. Three grass carps with similar body weight were put in each pond. The average body weights of grass carps put in each of the three ponds were about 0.5 kg, 1 kg, and 1.5 kg, respectively. The total body weights of fishes in each of the three ponds are 1.643 kg, 3.011 kg, and 5.270 kg, respectively. The fishes were raised for four days before the experiment started. During the five days of experiment, dead shrimps were fed with dead shrimps. The mean weight of dead shrimps used in the experiment is 5.3 g. Each day, the total body weight of dead shrimps that were ingested by fishes in each pond was calculated by subtracting the total body weight of dead shrimps remained in the pond from the total weight of dead shrimps put in the pond. The shrimp ingestion rate of fish is quantified by the daily ingestion rate (total body weight of ingested shrimps per day / total body weight of fishes).

To determine the rate of dead shrimp ingestion of African sharptooth catfish (*Clarias gariepinus*). Four cement ponds were set up with the same dimensions of 315cm × 315cm × 120cm, water volume of 5 $m^3$, and salinity of 3. One African sharptooth catfish with body weight of 0.262 kg, 0.496 kg, 0.731 kg, and 1.502 kg were put in each pond separately. The fishes were raised for four days before the experiment started. During the five days of experiment, fishes were fed with dead shrimps. The mean weight of dead shrimps used in the experiment is 6.2 g. Each day, the total body weight of dead shrimps that were ingested by fishes in each pond was calculated by subtracting the total body weight of dead shrimps remained in the pond from the total weight of dead shrimps put in the pond. The shrimp ingestion rate of fish is quantified by the daily ingestion rate (total body weight of ingested shrimps per day / body weight of fish).

To determine the rate of dead shrimp ingestion of red drum (*Sciaenops ocellatus*). Three cement ponds were set up with the same dimensions of 315cm × 315cm × 120cm, water volume of 5 $m^3$, and salinity of 5. One red drum with body weight of 0.590 kg, 0.654 kg, and 0.732 kg was put in each pond, separately. The fishes were raised for four days before the experiment started. During the five days of experiment, fishes were fed with dead shrimps. The mean weight of dead shrimps used in the experiment is 3.9 g. The daily total body weight of dead shrimps that were ingested by fishes in each pond was calculated by subtracting the total body weight of dead shrimps remained in the pond from the total weight of dead shrimps put in the pond. The shrimp ingestion rate of fish is quantified by the daily ingestion rate (total body weight of ingested shrimps per day / body weight of fish).

The ratio of healthy shrimp ingestion of fishes

To determine the rate of healthy shrimp ingestion of grass carp, three experimental groups and one control group were set up. Four cement ponds were set up with the same dimensions of 315cm × 315cm × 120cm, water volume of 5 $m^3$, and salinity of 5. Regarding the density of $7.5x10^5$-shrimps/$hm^2$ in shrimp farming production, 750 WSSV-free shrimps (average 5.3 g of body weight) were delivered into each concrete pond. Grass carps with body weight of 1.05 kg, 0.956 kg, and 1.013 kg were put in each experiment pond separately. No fish was put in the control pond. Every two days, 50% of the water in each pond was changed. Live shrimps remained in each pond was counted and weighted after 10 days of the experiment.

To determine the rate of healthy shrimp ingestion of African sharptooth catfish, one experimental group and one control group were set up. Two cement ponds were set up with the same dimensions of 315cm × 315cm × 120cm, water volume of 5 $m^3$, and salinity of 3. Regarding the density of $7.5x10^5$-shrimps/$hm^2$ in shrimp farming production, 750 WSSV-free shrimps (average 2.2 g of body weight) were delivered into each concrete pond. African sharptooth catfish with body weight of 1.05 kg was put in the experiment pond. No fish was put in the control pond. Every two days, 50% of the water in each pond was changed. Live shrimps remained in each pond was counted and weighted after 10 days of the experiment.

To determine the rate of healthy shrimp ingestion of red drum, three experimental groups and one control group were set up. Four cement ponds were set up with the same dimensions of 315cm × 315cm × 120cm, water volume of 5 $m^3$, and salinity of 5. Regarding the density of $7.5x10^5$-shrimps/$hm^2$ in shrimp farming production, 750 WSSV-free shrimps (average 2.7 g of body weight) were delivered into each concrete pond. Red drums with body weight of 0.519 kg, 0.554 kg, and 0.595 kg were put in the experiment pond. No fish was put in the control pond. Every two days, 50% of the water in each pond was changed. Live shrimps remained in each pond was counted and weighted after 10 days of the experiment.

The feeding selectivity of fish on dead, infected, and healthy shrimps

To determine the feeding selectivity of grass carp on dead, infected, and healthy shrimps, one aquarium was designed with the same dimension of 220 cm × 60 cm × 80 cm, water volume of 0.5 $m^3$, and salinity of 5. Grass carp with body weight of 1.58 kg was raised in the aquarium for four days before the experiment started. WSSV-infected shrimps in the latent period would die within one day as long as the individual converted to be diseased. Thus, it is impossible to use WSSV-infected shrimps directly in the experiment. The activity of shrimps reduced after the endopod and exopod removed, which are similar to the reduction of activity of diseased shrimps. Thus, the endopod and exopod removed shrimps were used to resemble WSSV-infected shrimps. Thirty pieces of healthy shrimps were put in the control aquarium. On day 1 of the experiment, 30 pieces each of dead, tailfin-cut, and healthy shrimps were put in the aquarium. The mean weight of shrimps used in the experiment is 3.5 g. Then the dead, infected (endopod and exopod removed), and healthy shrimps remained in the experimental aquarium were counted and weighted every 24 hours. New shrimps were added to ensure there are 30 pieces each of dead, infected (endopod and exopod removed), and healthy shrimps in the aquarium. The experiment continued for 9 days. The daily total body weight of shrimps that were ingested by fishes in each pond was calculated by subtracting the total body weight of shrimps remained in the pond from the total weight of shrimps put in the pond. The shrimp ingestion rate of fish is quantified by the daily ingestion rate (total body weight of ingested shrimps per day / body weight of fish).

To determine the feeding selectivity of African sharptooth catfish on dead, infected, and healthy shrimps, one aquarium was designed with the same dimension of 220 cm × 60 cm × 80 cm, water volume of 0.5 $m^3$, and salinity of 3. African sharptooth catfish with body weight of 1.03 kg was raised in the aquarium for four days before the experiment started. WSSV-infected shrimps in the latent period would die within one day as long as the individual converted to be diseased. Thus, it is impossible to use WSSV-infected shrimps directly in the experiment. The activity of shrimps reduced after the endopod and exopod removed, which are similar to the reduction of activity of diseased shrimps. Thirty pieces of healthy shrimps were put in the control aquarium. On day 1 of the experiment, 30 pieces each of dead, tailfin-cut, and healthy shrimps were put in the aquarium. The mean weight of shrimps used in the experiment is 8.4 g. Then the dead, infected (endopod and exopod removed), and healthy shrimps remained in the experimental aquarium were counted and weighted every 24 hours. New shrimps were added to ensure there are 30 pieces each of dead, infected (endopod and exopod removed), and healthy shrimps in the aquarium. The experiment continued for 9 days. The daily total body weight of shrimps that were ingested by fishes in each pond was calculated by subtracting the total body weight of shrimps remained in the pond from the total weight of shrimps put in the pond. The shrimp ingestion rate of fish is quantified by the daily ingestion rate (total body weight of ingested shrimps per day / body weight of fish).

Determine the bodyweight of grass carp suitable for control of shrimp WSS

To determine the body weight of grass carp for effective control of WSS, six groups were used: four experimental groups and two control groups. Each group consisted of three replicates, and each replicate was put in 600 pieces of 5 g shrimps. Four groups were separately set in one piece of grass carp of average body weight 0.3 kg, 0.5 kg, 1.0 kg, 1.5 kg, respectively, and separately put in 3 pieces of 5 g WSSV-infected shrimps. In two control groups, one was put in three WSSV infected shrimp without grass carp, and the other was put in a 1.1 kg grass carp without WSSV-infected shrimp. The number of live shrimps was recorded ten days of post-infection. In the groups with dead shrimps ten days post-infection, ten dead shrimps were selected for one-step WSSV PCR detection, showing positive for WSSV.

Determine the bodyweight of catfish suitable for control of shrimp WSS

To determine the body weight of catfish for effective control of WSS, six ponds (dimensions of 220 cm × 60 cm × 80 cm, water volume of 0.5 $m^3$) were used: four experimental ponds and two control ponds. Each pond was put in 600 pieces of 1.5 g shrimps, of which 20% are WSSV carriers. Four groups were separately set in one piece of African sharptooth catfish of average body weight 0.25 kg, 0.5 kg, 0.75 kg, 1.5 kg, respectively, and separately put in 3 pieces of 1.5 g WSSV-infected shrimps. In two control groups, one was put in three WSSV infected shrimp without African sharptooth catfish, and the other was put in a 0.76 kg African sharptooth catfish without WSSV-infected shrimp. The number of live shrimps was recorded ten days of post-infection. In the groups with dead shrimps ten days post-infection, ten dead shrimps were selected for one-step WSSV PCR detection, showing positive for WSSV.

The capacity of grass carp for controlling WSS

To determine the capacity of grass carp for controlling WSS, the number of WSSV-infected shrimps that could be cleared by one 1kg-grass-carp was evaluated. Shrimps of different specifications (1.3 ± 0.1 g, 2.5 ± 0.2 g, 5.0 ± 0.3 g, or 7.8 ± 0.5 g of body weight, respectively) were separately used in four ponds, in which WSSV acutely infected shrimps were put in. Artificially WSSV acutely infected shrimps were obtained as follows: In one pond, healthy shrimps were starved from 3 days and then fed twice for 24 hours with 20% feed of WSSV-infected dead shrimp body weight, i.e., 20g dead shrimps for 100g healthy shrimps. Such a 20% ratio of dead shrimp versus healthy shrimps (by body weight) could ensure to obtain the WSSV acute infection shrimps. Ten shrimps were used to detect the presence of WSSV via the one-step PCR assay. These WSSV acutely infected shrimps were used for the following experiments.

In 1.3 ± 0.1 g group, 750 healthy shrimps (WSSV-free) were placed into each of the nine cement ponds (size of 10 $m^2$, water volume of 5 $m^3$). The 750 shrimps/10$m^2$ was calculated from the farm culture density of $7.5x10^5$-shrimps/$hm^2$. One piece of average 1 kg grass carp was put in seven ponds containing 3, 6, 9, 12, 15, 18, or 21 pieces, respectively, of about 1.3 g WSSV acutely infected shrimps. The other two ponds were used as controls: One pond was not put in grass carp but put in three WSSV acutely infected shrimps as a positive control; the other pond was put in neither grass carp, nor WSSV acutely infected shrimps, serving as a negative control. Shrimps were fed with 2% artificial feed of body weight. And 50% of the water was changed every day. The experiments were concluded by counting the number of live shrimps 15 days post-infection.

In 2.5 ± 0.2 g group, the number of WSSV acutely infected shrimps was adjusted per the results of the above 1.3 ± 0.1 g group. Specifically, 10, 20, 30, 40, 50, 60, or 70 pieces of WSSV acutely infected shrimps were respectively put in seven experimental ponds. The positive control pond was put in 10 WSSV acutely infected shrimp while the negative control pond was set as the same as in the 1.3 ± 0.1 g set of experiments. All other culture conditions and data collection were the same as above.

In 5.0 ± 0.3 g group, the number of WSSV acutely infected shrimps was adjusted per the results of the above 2.5 ± 0.2 g group. Specifically, 50, 70, 90, 110, 120, 130, or 140 pieces of WSSV acutely infected shrimps were respectively put in each of the seven experimental ponds. The positive control pond was placed in 50 WSSV acutely infected shrimps while the negative control pond was set as the same as in the above 1.3 ± 0.1 g group of experiments. All other culture conditions and data collection were the same as above. In a 7.8 ± 0.5 g group, the number of WSSV acutely infected shrimps was adjusted per the results of the above 5.0 ± 0.3 g group. Six cement ponds (size of 10 $m^2$, water volume of 5 $m^3$) were respectively put in 750 pieces of matching body weight of healthy shrimps (WSSV-free). Specifically, 30, 40, 50, or 60 pieces of WSSV acutely infected shrimps, were respectively put in each of 4 experimental ponds (size of 10 $m^2$, water volume of 5 $m^3$), followed by putting in one piece of 1.0 ± 0.05 kg grass carp. The positive control pond was put in 30 WSSV acutely infected shrimps while the negative control pond was set as the same as in the above 1.3 ± 0.1 g group of experiments. All other culture conditions and data collection were the same as above.

Based on the relationship of healthy shrimp, infected shrimp, dead shrimp, and fish, Model 3 was established for illustrating the control of WSS by fish.

Determine the number of grass carps required for prevention of WSS in shrimp production

To validate the above laboratory experiments in a field, the number of grass carps required for the prevention of shrimp WSS in shrimp farming practice was first determined in 2010 by using 40 farm-size ponds (Area of 0.34 ± 0.04 $hm^2$/pond) in Pinggang Aquaculture Base, Yangjiang, Guangdong Province, China. These 40 ponds were divided into eight groups; each group consisted of 5 ponds; The amount of 675,000-shrimp/$hm^2$ of Pacific white shrimp (body length 0.8~ 1.0 cm) was released into ponds. The #1 group of ponds was not put in grass carps. The #2, #3, #4, #5, #6, #7, #8 groups of ponds were respectively put in 45, 150, 225, 300, 450, 600, 750-grass carps/$hm^2$, with a bodyweight of 1.0 kg~1.25 kg. These 40 ponds were managed by using the same farming method. If the WSS outbreak occurred, shrimps were harvested; if not, shrimps were harvested at 110 days, and yields were measured.

Validation of technology in the application of grass carps for WSS control

Based on the above experimental results, the standard specification and quantity of grass carps required for the prevention of shrimp WSS as follows: a bodyweight of 1.0 kg~1.25 kg and 300~450-grass-carps/$hm^2$. In 2011, this standard specification was verified in Guan-Li-Da Company, Maoming, Guangdong Province, China. Forty-six farm ponds were used with total area 17.33 $hm^2$ and each pond of 0.377 $hm^2$. The 46 ponds were divided into zone A and zone B: Zone A consisted of 18 ponds of total area 6.03 $hm^2$ while zone B consisted of 28 ponds of total area 11.30 $hm^2$. Zone A ponds were put in 900,000-shrimp/$hm^2$ and 20 days after stocking followed up with releasing 317 ~ 450-grass-carp/$hm^2$ of average body weight of 1.0 kg. Zone B was put in only 900,000-shrimp/$hm^2$ without grass carps. In 2012, Zones A and B were switched, i.e., A without grass carps while B with grass carps. If WSS outbreak occurred, shrimps were harvested; if not, shrimps were harvested at 110 days, and yields were measured.

Validation of technology in the application of catfish for WSS control

In 2011 and 2012, this standard specification was verified in Fushun shrimp farms, Qinzhou, Guangxi Province, China. Ninety-five farm ponds were used with a total area of 88.2 $hm^2$. In 2011, the 95 ponds were divided into zone A and zone B: Zone A consisted of 38 ponds of total area 21.2 $hm^2$ while zone B consisted of 57 ponds of total area 67.0 $hm^2$. Zone A ponds were put in 750,000-shrimps/$hm^2$ (body length of 0.8~1.0cm) and 10 days after stocking followed up with releasing 525 ~ 750-catfish/$hm^2$ of average body weight of 0.5 kg. Zone B was put in only 750,000-shrimps/$hm^2$ without catfish. In 2012, zone A was the same as in 2011 (shrimp ponds with catfish). However, zone B was divided into zone B1 and zone B2: Zone B1 consisted of 25 ponds of total area 27.0 $hm^2$ while zone B2 consisted of 32 ponds of total area 40.0 $hm^2$. Zone B1 ponds were set the same as the zone A in 2011 while zone B2 was set the same as the zone B in 2011. If WSS outbreak occurred, shrimps were harvested; if not, shrimps were harvested at 110 days, and yields were measured.

Validation of using fishes for WSS control in the actual shrimp production

From 2013 to 2019, we tested the effectiveness of using fish for controlling WSS in actual shrimp production at the Guan-Li-Da Company, Maoming, Guangdong Province, China. In 2013, shrimps were co-cultured with African sharptooth catfish of body weight ranges from 0.25~0.5 kg in 13 ponds. The density of shrimps in these ponds ranges from 878,788-shrimps/$hm^2$ to 1,230,769-shrimps/$hm^2$. And shrimps were co-cultured with grass carp of body weight ranges from 0.65-0.8 kg and African sharptooth catfish of body weight ranges from 0.25~0.5 kg in 10 ponds. The density of shrimps in these ponds ranges from 909,091-shrimps/$hm^2$ to 2,000,000-shrimps/$hm^2$. Additionally, shrimps were cultured without fish in 11 ponds. The density of shrimps in these ponds ranges from 878,788-shrimps/$hm^2$ to 969,697-shrimps/$hm^2$. If WSS outbreak occurred, shrimps were harvested; if not, shrimps were harvested at 110 days, and yields were measured.

In 2014, shrimps were co-cultured with grass carp of average body weight of 0.75 kg in 8 ponds. The density of shrimps in these ponds ranges from 161,111-shrimps/hm$^2$ to 181,818-shrimps/hm$^2$. And shrimps were co-cultured with grass carp of average body weight of 0.75 kg and African sharptooth catfish of average body weight of 0.35 kg in 12 ponds. The density of shrimps in these ponds ranges from 145,000-shrimps/hm$^2$ to 181,818-shrimps/hm$^2$. Additionally, shrimps were cultured without fish in 5 ponds. The density of shrimps in these ponds was 181,818-shrimps/hm$^2$. If WSS outbreak occurred, shrimps were harvested; if not, shrimps were harvested at 110 days, and yields were measured.

In 2015, shrimps were co-cultured with grass carp of average body weight of 0.75 kg in 19 ponds. The density of shrimps in these ponds ranges from 746,269-shrimps/hm$^2$ to 1,538,462-shrimps/hm$^2$. Additionally, shrimps were cultured without fish in 10 ponds. The density of shrimps in these ponds ranges from 750,000-shrimps/hm$^2$ to 909,091-shrimps/hm$^2$. If WSS outbreak occurred, shrimps were harvested; if not, shrimps were harvested at 110 days, and yields were measured.

In 2016, shrimps were co-cultured with grass carp of average body weight of 0.75 kg in 19 ponds. The density of shrimps in these ponds ranges from 488,372-shrimps/hm$^2$ to 636,364-shrimps/hm$^2$. Additionally, shrimps were cultured without fish in 8 ponds. The density of shrimps in these ponds ranges from 543,478-shrimps/hm$^2$ to 636,364-shrimps/hm$^2$. If WSS outbreak occurred, shrimps were harvested; if not, shrimps were harvested at 110 days, and yields were measured.

In 2017, shrimps were co-cultured with grass carp of average body weight of 0.75 kg in 6 ponds. The density of shrimps in these ponds was 961,538-shrimps/hm$^2$. And shrimps were co-cultured with grass carp of average body weight of 0.75 kg and African sharptooth catfish of average body weight of 0.15 kg in 12 ponds. The density of shrimps in these ponds ranges from 848,485-shrimps/hm$^2$ to 909,091-shrimps/hm$^2$. Additionally, shrimps were cultured without fish in 9 ponds. The density of shrimps in these ponds ranges from 848,485-shrimps/hm$^2$ to 961,538-shrimps/hm$^2$. If WSS outbreak occurred, shrimps were harvested; if not, shrimps were harvested at 110 days, and yields were measured.

In 2018, shrimps were co-cultured with grass carp of average body weight of 0.75 kg in 22 ponds. The density of shrimps in these ponds ranges from 378,788-shrimps/hm$^2$ to 869,565-shrimps/hm$^2$. Additionally, shrimps were cultured without fish in 9 ponds. The density of shrimps in these ponds ranges from 695,652-shrimps/hm$^2$ to 861,111-shrimps/hm$^2$. If WSS outbreak occurred, shrimps were harvested; if not, shrimps were harvested at 110 days, and yields were measured.

In 2019, shrimps were co-cultured with grass carp of average body weight of 0.75 kg in 30 ponds. The density of shrimps in these ponds ranges from 652,174-shrimps/hm$^2$ to 1,000,000-shrimps/hm$^2$. Additionally, shrimps were cultured without fish in 10 ponds. The density of shrimps in these ponds ranges from 500,000-shrimps/hm$^2$ to 1,000,000-shrimps/hm$^2$. If WSS outbreak occurred, shrimps were harvested; if not, shrimps were harvested at 110 days, and yields were measured.

**Supplementary Text**

Mathematical model 1 - The relationship among the body weight of the initial WSSV-infected shrimp, number of deaths, and death time distribution.

The experimental data show the time course of death for the infected shrimps satisfies the Laplacian distribution (table S2-4). The relationship of the body weight of the initial infected shrimp number of deaths, and death time distribution could be expressed by a mathematical model and the establishment of the mathematical model was shown below.

Suppose that one dead shrimp could infect *n* healthy shrimps at the same day. These *n* infected shrimps do not die simultaneously but in different days (time course). The value of *n* is related to the weight of the dead shrimps - Large dead shrimp can infect more healthy shrimps of the same body weight. Our experimental results (table S2-4) show the death time course for these *n* infected shrimps satisfies the Laplacian distribution, as follows:

$$p(t) = \begin{cases} b\exp\left(-\frac{|t-a|}{c_1}\right), & t \le a \\ b\exp\left(-\frac{|t-a|}{c_2}\right), & t > a \end{cases} \quad (1)$$

where *a* is the peak time of number of dead shrimps, *b* is the maximal death percentage, $c_1$ is related to the mortality increases of the infected shrimps, $c_2$ is related to the mortality decreases of the infected shrimps, *p*(*t*) is the percentage of infected shrimps that die at time *t*. The open bracket "{" in formula (1) means the function is represented by two parallel expressions as described previously.

Based on the table S2-4, we can determine the value of *a*, *b*, $c_1$ and $c_2$ by the least square estimation (LSE) method. As different weight corresponds to different distribution of death time, we can compute the relationship of weight of death shrimps with corresponding *a*, *b*, $c_1$ and $c_2$ (table S20).

We found the relationship of *w* with *a*, or *b*, or $c_1$ or $c_2$ is quadratic (Equation 2), with the data in table S20, we have

$$\begin{aligned} a &= -0.0918w^2 + 0.8772w + 3.3449 \\ b &= 0.0029w^2 - 0.0369w + 0.5849 \\ c_1 &= -0.0186w^2 + 0.1739w + 0.7063 \\ c_2 &= 0.0002w^2 + 0.0108w + 1.0827 \end{aligned} \tag{2}$$

Using Model 1, we can predict the effects of different body weights of WSSV infected dead shrimps through the ingestion pathway of WSSV-infected dead shrimps on the WSSV transmission rate.

<u>Mathematical Model 2 – the dynamic changes of healthy, infected, and dead shrimps during WSSV transmission</u>

We derived and established Model 2 to simulate the WSS transmission dynamics in cultured shrimps. Using Model 2, we predicted the dynamic exchanges of three states (healthy, infected, and dead shrimps) in cultured shrimps as influenced by WSS epidemic with the following:

Now we can develop a model for the spread and break out of WSS. For any given weight *w* of shrimps, let $s_h(t)$, $s_i(t)$ and $s_d(t)$ be the number of healthy shrimps, infected shrimps and dead shrimps respectively at time *t*. Let $I(t)$, $d(t)$ be the number of daily infected shrimps, daily dead shrimps, respectively, at time *t*.

According to infection process, the decrement of healthy shrimps is caused by their infection, therefore we have $\frac{ds_h}{dt} = -I(t)$. The quantity change of infected shrimps includes the infection of healthy shrimps and the death of infected shrimps, we have $\frac{ds_i}{dt} = I(t) - d(t)$. The increment of dead shrimps is caused by the death of the infected shrimps; thus we have $\frac{ds_d}{dt} = d(t)$. We obtain the following system of ordinary differential equations:

$$\frac{ds_h}{dt} = -I(t)$$
$$\frac{ds_i}{dt} = I(t) - d(t) \qquad (3)$$
$$\frac{ds_d}{dt} = d(t)$$

where $s_h(0)= s_{h0}$, $s_i(0)= s_{i0}$, $s_d(0)= s_{d0}$ are as the initial value, at $t$=0.

In the above system of ordinary differential equations, quantity $I(t)$ can be expressed as follows

$$I(t) = \min\{ns_d(t), s_h(t) - \alpha s_{h0}\} \qquad (4)$$

$d(t)$ can be expressed as

$$d(t) = \int_0^t \min\{ns_d(t-\tau), s_h(t-\tau) - \alpha s_{h0}\} p(\tau) d\tau \qquad (5)$$

where $n$ is the number of healthy shrimps infected by one dead shrimp the first day. $p(\tau)$ is the death percentage of the $n$ infected shrimps at $\tau$ days, $T$ is the longest survival time of infected.

Now we explain how to set up the formulas $I(t)$ and $d(t)$. In the expression of $I(t)$ , $ns_d(t)$ is the number of daily infected shrimps at time $t$. But as the number of healthy shrimps decreases, there may not be as many as $ns_d(t)$ healthy shrimps to be infected. Therefore, $I(t)$ is the minimum of $ns_d(t)$ and $s_h(t) - \alpha s_{h0}$ , where $\alpha$ ( $0 < \alpha < 1$) represents the percentage of healthy shrimps that may have resistance to viruses, $d(t)$ is the number of shrimps infected from 0 to $t$ die at time $t$. We use this integral to express the number of shrimps die at time $t$.

To evaluate the performance of the model 2, we compare the simulated scenario and the biological experimental settings. Our experiments show the quantity change of dead shrimps and live shrimps with respect to time, which is consistent with the result of simulation (figure S5).

Mathematical Model 3 - Use Fish to Control WSS

We established Model 3 for the prevention and control of WSS using fish. In Model 3, two parameters need to be determined before this model can be applied for evaluating the fish's capability of WSS prevention and control. The two parameters are, (1) fish-feeding quantity of dead shrimps, and (2) fish-feeding ratio of dead shrimps with healthy shrimps. We obtained 1 kg grass carp's feeding quantity of different body weights of shrimps and the feeding selectivity through experiments. The mathematical reasoning of Model 3 is as follows:

To block the transmission of WSS, we apply fish to eat dead shrimps and infected shrimps. Let $e_h(t)$, $e_i(t)$ and $e_d(t)$, respectively be the number of healthy shrimps, infected shrimps and dead shrimps eaten by fish daily at time $t$, $f(t)$ is the number of fish.

The decrement of healthy shrimps is related to the number of infected healthy shrimp and the number of shrimps eaten by fish, as expressed in $\frac{ds_h}{dt} = -I(t) - e_h(t)$. Similarly, the dynamics of the infected shrimps is related to the number of infected healthy shrimps, the number of death of infected shrimps, and the number of infected shrimps eaten by fish, as expressed in $\frac{ds_i}{dt} = I(t) - d(t) - e_i(t)$. The dynamics of dead shrimps is related to the number of deaths of infected shrimps, and eaten by fish, as expressed in $\frac{ds_d}{dt} = d(t) - e_d(t)$. Combining the above formulae, we can write the model as follows:

$$\begin{aligned} \frac{ds_h}{dt} &= -I(t) - e_h(t) \\ \frac{ds_i}{dt} &= I(t) - d(t) - e_i(t) \\ \frac{ds_d}{dt} &= d(t) - e_d(t) \end{aligned} \tag{6}$$

where $s_h(0)=s_{h0}$, $s_i(0)=s_{i0}$, $s_d(0)=s_{d0}$ are as the initial value at $t$=0. In the above model, $I(t)$, $d(t)$, $e_h(t)$, $e_i(t)$ and $e_d(t)$ are respectively given as follows:

$$
\begin{aligned}
I(t) &= \min\{ns_d(t),\, s_h(t) - \alpha s_{h0}\} \\
d(t) &= \int_0^t \min\{ns_d(t-\tau),\, s_h(t-\tau) - \alpha s_{h0}\} p(\tau) \exp\left\{\int_{t-\tau}^{t} \ln r(u) du\right\} d\tau \\
e_d(t) &= \min\{f(t)\cdot m \cdot \beta,\ \ s_d(t) + d(t)\} \\
e_i(t) &= \min\left\{(f(t)\cdot m - e_d(t))\frac{s_i(t) + I(t) - d(t)}{s_i(t) + s_h(t) - d(t)},\ \ s_i(t) + I(t) - d(t)\right\} \\
e_h(t) &= \min\{f(t)\cdot m - e_d(t) - e_i(t),\ \ s_h(t) - I(t)\} \\
r(t) &= 1 - \frac{e_i(t)}{s_i(t) + I(t) - d(t)}
\end{aligned} \tag{7}
$$

where, $I(t)$ is the same as in equation (4); for $d(t)$, different from equation 5 is that we add an exponential item $\exp\left\{\int_{t-\tau}^{t} \ln r(u)du\right\}$ to account for the infected shrimps that may be eaten by fish during the past t days. As for $e_d(t)$ shown in equation (6), $m$ is for that each fish eats $m$ shrimps while $\beta$ accounts for percentage of dead shrimps in $m$ shrimps. In $e_i(t)$, we introduce $\frac{s_i(t) + I(t) - d(t)}{s_i(t) + s_h(t) - d(t)}$ for the percentage of infected shrimps in live shrimps. $e_h(t)$ accounts for the number of healthy shrimps eaten by fish. $r(t)$ represents the percentage of infected shrimps not being eaten by fish. We performed the effects of 1 kg grass carps on shrimps with four different body weights. The simulated data agreed with the experimental results agreed with (Figure 2B).

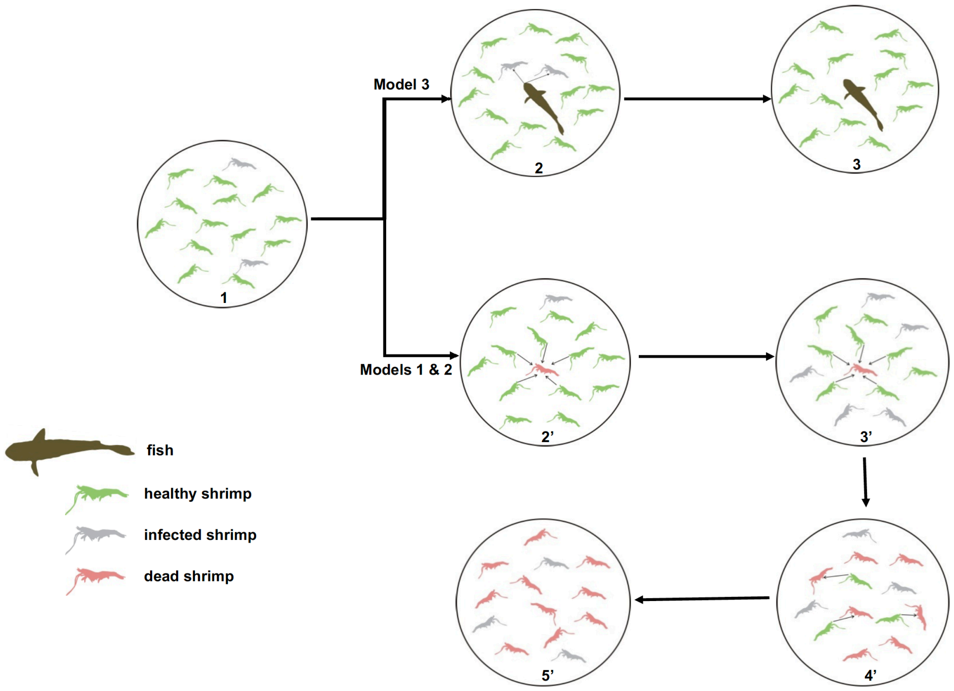


**Fig. S1.**

**The schematic diagram for controlling WSSV outbreaks by fish in shrimp population.**

<u>Models 1 and 2: WSS transmission process through healthy shrimps ingesting infected dead shrimps.</u>

1) Virus-carrier shrimps are converted to be diseased shrimps upon environmental stress, initiating the WSS transmission in the shrimp population.

2) Diseased shrimps die; healthy shrimps ingest the infected dead shrimps.

3) The healthy shrimps that ingested infected dead shrimps become diseased shrimps and die. Other healthy shrimps continue to ingest these infected dead shrimps.

4) The cycle of the infection (from #2 to #4) continues in the shrimp population.

5) The cycle of infection drives the WSS outbreak, leading to a shrimp population collapses in a pond.

<u>Model 3: Preventing WSSV outbreak in shrimp population by co-culturing fishes.</u>

1) Virus-carrier shrimps are converted to be infected shrimps upon environmental stress, initiating the WSS transmission in a shrimp population.

2) Fishes ingest diseased shrimps and infected dead shrimps.

3) Fishes remove diseased and infected dead shrimps, keeping the shrimp population healthy

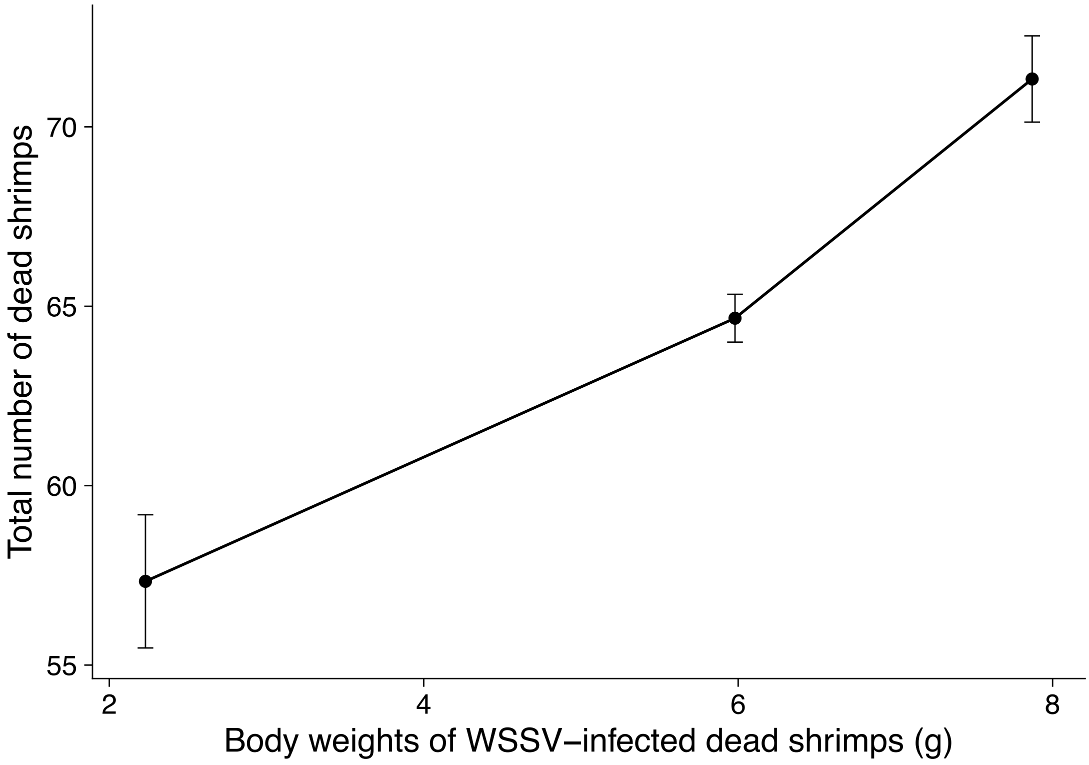


**Fig. S2.**

**Numbers of healthy shrimps died from WSSV infection in aquariums adding one piece of WSSV-infected dead shrimp of body weight of 1.98 g, 6.13 g, and 7.95 g, respectively**. Means and standard errors are shown. With the initial WSSV-infected dead shrimps become heavier, the basic reproduction number ($R_0$) of WSSV increases.

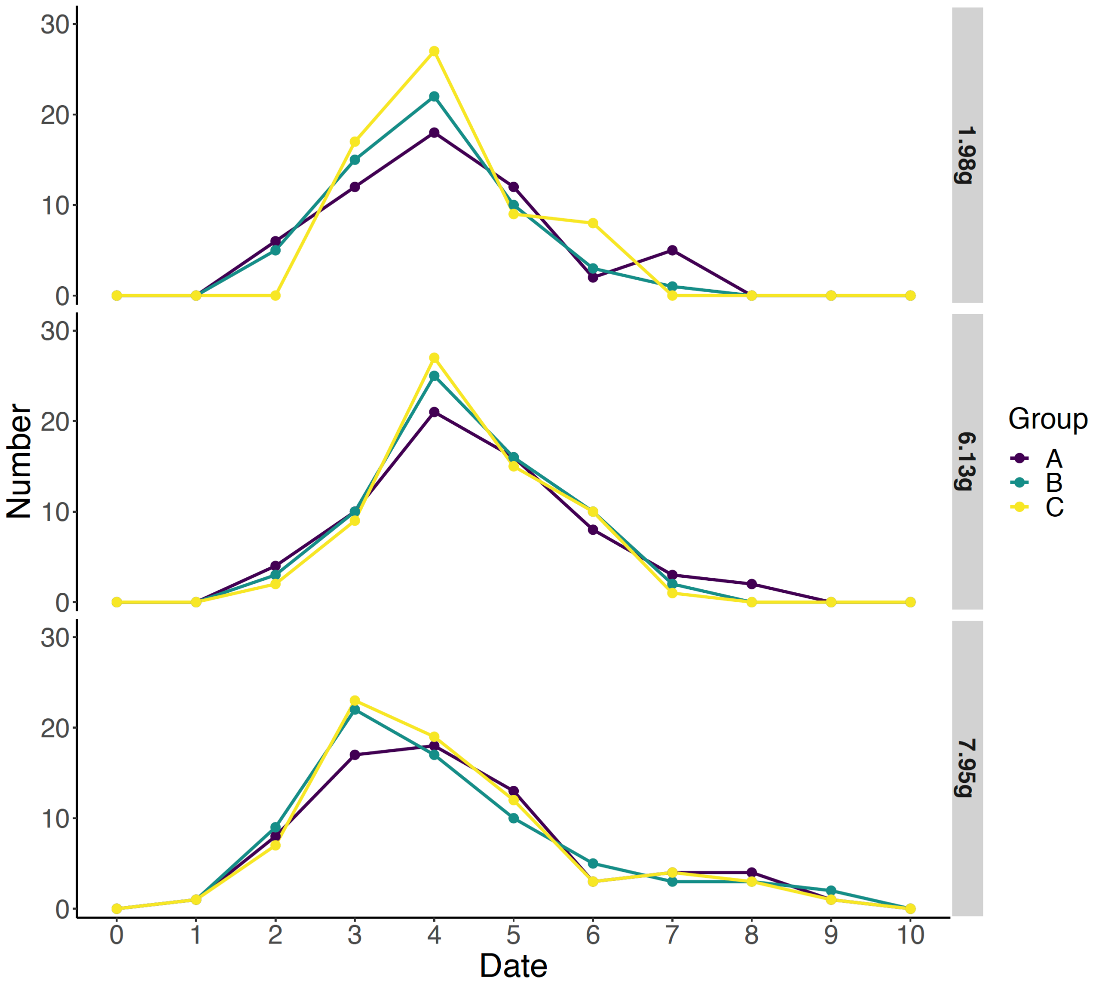


**Fig. S3.**

**Daily mortality of healthy shrimps in aquariums with one piece of WSSV-infected dead shrimp of body weights of 1.98 g, 6.13 g, and 7.95 g, respectively.** Time to death was consistent across the three groups of body weight for WSSV-infected shrimps, with the majority on the third to sixth day and the peak number of deaths on the fourth and fifth days.

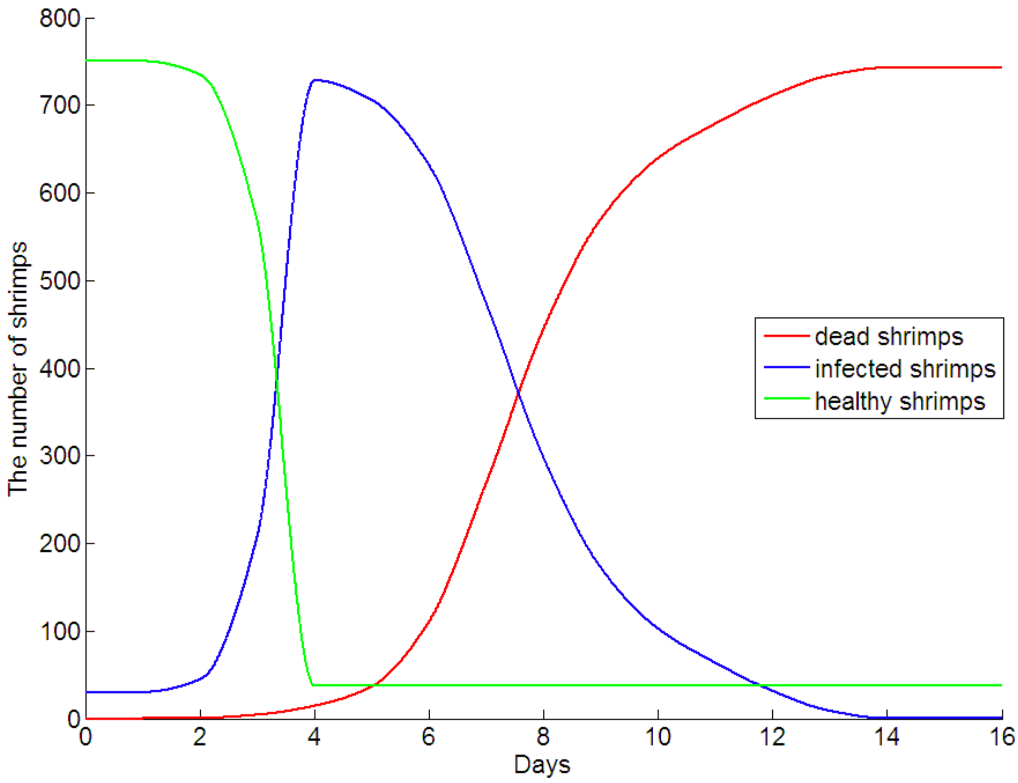


**Fig. S4.**

**The changes of number of healthy, WSSV-infected, and dead shrimps over time.** The numbers of healthy shrimps (green), infected shrimps (blue) and dead shrimps (red) were derived from Model 2. The number of infected healthy shrimps drastically increased after 2 days of WSSV infection. The number of infected shrimps began to decrease after the 4th day as infected shrimps became dead shrimp, which led to the increasing of dead shrimps. Deaths caused by WSSV infection rose sharply after 6 days. Finally, all the shrimps became dead shrimps except for a small proportion of shrimps that might have resistance to WSSV.

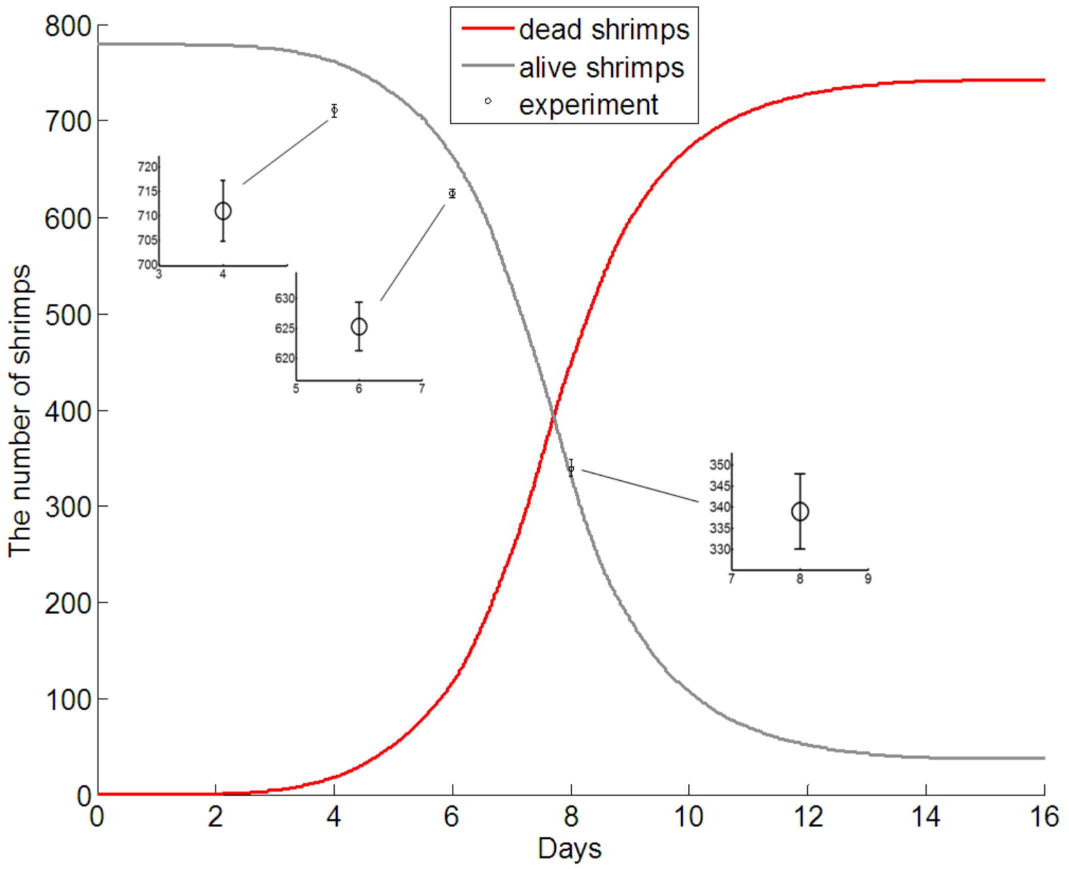


**Fig. S5.**

**The quantity of dead shrimps (red) and live shrimps (grey) (include healthy shrimps and infected shrimps) concerning time.** The red curve and the grey curve are the results of the simulation. The three small open circles (dots) with error bars were the numbers of live shrimps on the fourth, sixth, eighth day from the artificial infection experiments. This data set is derived from three repeats of independent experiments, which is expressed as mean ± SD. The experimental results were consistent with the results of mathematical modeling.

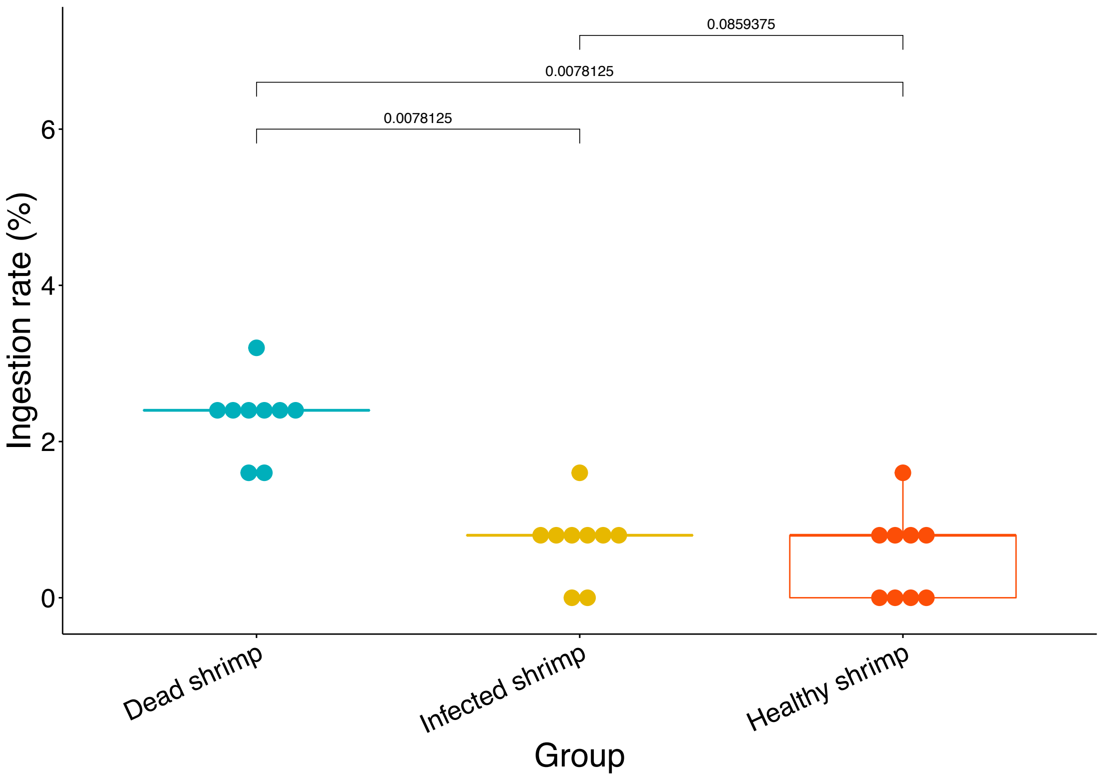


**Fig. S6.**

**Feeding selectivity of African sharptooth catfish on dead, infected (endopod and exopod removed), and healthy shrimps.** The WSSV-infected shrimps would die within one day. In addition, the activity of shrimps reduced after the endopod and exopod removed, which are similar to the reduction of activity of WSSV-infected shrimps. Thus, the endopod and exopod removed shrimps were used to resemble WSSV-infected shrimps. One piece of grass carp was co-cultured with dead, infected (endopod and exopod removed), and healthy shrimps in one pond. Different kinds of shrimps that were ingested by fish were counted each day. The shrimp ingestion rate of fish is quantified by the daily ingestion rate (weight of shrimps ingested by fish per day / fish body weight). The body weight of grass carp is 1.05 kg. The mean weight of shrimps used in the experiment is 8.2 g. *P*-values (permutation test, paired) were labeled. African sharptooth catfish ingested significantly more dead shrimps than infected (endopod and exopod removed) and healthy shrimps.

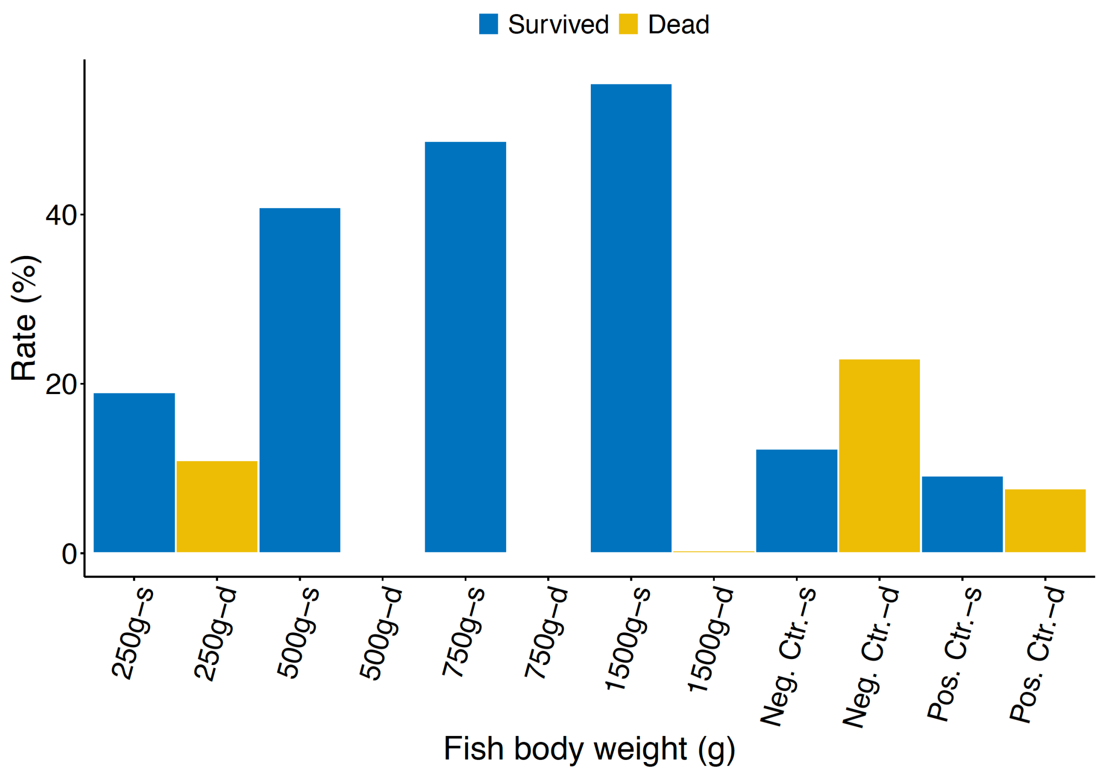


**Fig. S7.**

**The effect of the different body weights of African sharptooth catfish on the prevention and control of shrimp viral disease spreads.** Four experimental groups were established for catfish, in which 600 pieces of 1.5g shrimps were cultured with 3 WSSV-infected shrimps of same body weight as well as one catfish of different body weights. As it might have WSSV carriers in shrimp postlarva during shrimp production, we replaced 20% of healthy shrimps as WSSV carriers. After 14 days in culture, 19%, 40.83%, 48.67%, 55.5% of shrimps were survived in the experimental groups with one catfish weighted 0.25kg, 0.5kg, 0.75kg, 1.5kg, respectively. In addition, 11% of shrimps were dead and remained in the pond (not removed by fishes) in which the co-cultured catfish weighted 0.25kg. Whereas, almost all dead shrimps were removed by fishes in other experimental groups. This suggests the minimum body weight of co-cultured catfish to control WSS outbreak is 0.5kg. Furthermore, co-culturing of fish can control the outbreak of WSS even there are WSSV-carriers in shrimp postlarva.

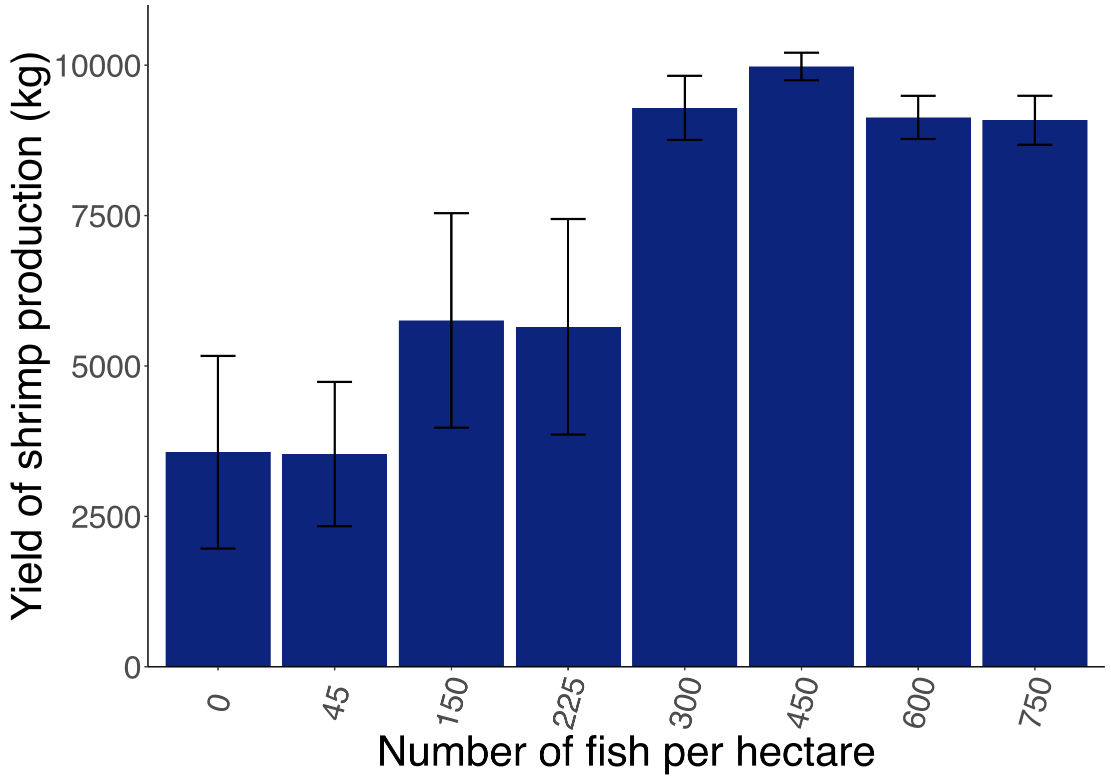


**Fig. S8.**

**The relationship of the number of co-culturing grass carp and the yield of shrimp production.** More than 300 grass carps of about 1 kg per hectare can completely control the outbreaks of WSS. Thus, the ponds that co-culturing more than 300 grass carps have substantial increment of the yield than the ponds that co-culturing less than 225 grass carps.

**Table S1.**

Numbers of healthy shrimps died from WSSV infection in aquariums adding one piece of WSSV-infected dead shrimp of body weights of 1.98 g, 6.13 g, and 7.95 g, respectively

| | 1.98 g | 6.13 g | 7.95 g |
|---|---|---|---|
| Group A | 55 | 64 | 69 |
| Group B | 56 | 66 | 72 |
| Group C | 61 | 64 | 73 |
| Mean | 57.3 | 64.7 | 71.3 |
| Negative Control | 1 | 0 | 2 |

**Table S2.**

Daily mortality of healthy shrimps in aquariums with one piece of WSSV-infected dead shrimp weighted 1.98 g

| Date | Aquarium A | Aquarium B | Aquarium C | Negative Control |
|---|---|---|---|---|
| **0** | 0 | 0 | 0 | 0 |
| **1** | 0 | 0 | 0 | 0 |
| **2** | 6 | 5 | 0 | 0 |
| **3** | 12 | 15 | 17 | 1 |
| **4** | 18 | 22 | 27 | 0 |
| **5** | 12 | 10 | 9 | 0 |
| **6** | 2 | 3 | 8 | 0 |
| **7** | 5 | 1 | 0 | 0 |
| **8** | 0 | 0 | 0 | 0 |
| **9** | 0 | 0 | 0 | 0 |
| **10** | 0 | 0 | 0 | 0 |

**Table S3.**

Daily mortality of healthy shrimps in aquariums with one piece of WSSV-infected dead shrimp weighted 6.13 g

| Date | Aquarium A | Aquarium B | Aquarium C | Negative Control |
|---|---|---|---|---|
| **0** | 0 | 0 | 0 | 0 |
| **1** | 0 | 0 | 0 | 0 |
| **2** | 4 | 3 | 2 | 0 |
| **3** | 10 | 10 | 9 | 0 |
| **4** | 21 | 25 | 27 | 0 |
| **5** | 16 | 16 | 15 | 0 |
| **6** | 8 | 10 | 10 | 0 |
| **7** | 3 | 2 | 1 | 0 |
| **8** | 2 | 0 | 0 | 0 |
| **9** | 0 | 0 | 0 | 0 |
| **10** | 0 | 0 | 0 | 0 |
| **11** | 0 | 0 | 0 | 0 |

**Table S4.**

Daily mortality of healthy shrimps in aquariums with one piece of WSSV-infected dead shrimp weighted 7.95 g

| Date | Aquarium A | Aquarium B | Aquarium C | Negative Control |
|---|---|---|---|---|
| **0** | 0 | 0 | 0 | 0 |
| **1** | 1 | 1 | 1 | 0 |
| **2** | 8 | 9 | 7 | 0 |
| **3** | 17 | 22 | 23 | 0 |
| **4** | 18 | 17 | 19 | 0 |
| **5** | 13 | 10 | 12 | 0 |
| **6** | 3 | 5 | 3 | 1 |
| **7** | 4 | 3 | 4 | 0 |
| **8** | 4 | 3 | 3 | 0 |
| **9** | 1 | 2 | 1 | 1 |
| **10** | 0 | 0 | 0 | 0 |
| **11** | 0 | 0 | 0 | 0 |
| **12** | 0 | 0 | 0 | 0 |

**Table S5.**

Daily dead shrimp ingestion of brown-marbled grouper (*Epinephelus fuscoguttatus*) with different body weights

| Group | 0.03 kg | 0.15 kg | 0.30 kg |
|---|---|---|---|
| | Pond A (0.098 kg) | Pond B (0.501 kg) | Pond C (0.999 kg) |
| Day 1 | 6.02% | 0 | 1.77% |
| Day 2 | 6.02% | 1.18% | 0.59% |
| Day 3 | 0 | 0 | 0 |
| Day 4 | 6.02% | 2.36% | 0.59% |
| Day 5 | 0 | 1.18% | 1.77% |
| Mean | 3.61% | 0.94% | 0.94% |

**Note**: The shrimp ingestion rate of fish is quantified by the daily ingestion rate (weight of shrimps ingested by fish per day / fish body weight). As the brown-marbled grouper did not eat any shrimp if only one fish was putted in the pond, three fishes were putted in each pond. The total body weight of fishes in each pond is listed below the pond name. The mean weight of dead shrimps used in the experiment is 5.9 g.

**Table S6.**

Daily dead shrimp ingestion of grass carp (*Ctenopharyngodon idellus*) with different body weights

| Group | 0.5 kg | 1 kg | 1.5 kg |
|---|---|---|---|
| | Pond A (1.643 kg) | Pond B (3.011 kg) | Pond C (5.27 kg) |
| Day 1 | 9.53% | 8.53% | 7.17% |
| Day 2 | 9.72% | 8.16% | 7.30% |
| Day 3 | 9.13% | 8.88% | 7.10% |
| Day 4 | 8.88% | 7.84% | 7.15% |
| Day 5 | 9.09% | 8.14% | 7.28% |
| Mean | 9.27% | 8.31% | 7.20% |

**Note:** The shrimp ingestion rate of fish is quantified by the daily ingestion rate (weight of ingested shrimps per day / fish weight). As the grass carp did not eat any shrimp if only one fish was putted in the pond, three fishes were putted in each pond. The total body weight of fishes in each pond is listed below the pond name. The mean weight of dead shrimps used in the experiment is 5.3 g.

**Table S7.**

Daily dead shrimp ingestion of African sharptooth catfish (*Clarias gariepinus*) with different body weights

| | Pond A (0.262 kg) | Pond B (0.496 kg) | Pond C (0.731 kg) | Pond D (1.502 kg) |
|---|---|---|---|---|
| Day 1 | 7.2% | 6% | 4.8% | 4.4% |
| Day 2 | 9.6% | 6% | 4.8% | 4% |
| Day 3 | 7.2% | 6% | 4% | 4.4% |
| Day 4 | 7.2% | 6% | 5.6% | 4.8% |
| Day 5 | 7.2% | 4.8% | 4.8% | 4.4% |
| Mean | 7.68% | 5.76% | 4.8% | 4.4% |

**Note**: The body weight of fishes in each pond is listed below the pond name. The shrimp ingestion rate of fish is quantified by the daily ingestion rate (weight of shrimps ingested by fish per day / fish body weight). The mean weight of dead shrimps used in the experiment is 6.2 g.

**Table S8.**

Daily dead shrimp ingestion of red drum (*Sciaenops ocellatus*) with different body weights

| | Pond C (0.59 kg) | Pond B (0.654 kg) | Pond A (0.732 kg) |
|---|---|---|---|
| Day 1 | 9.33% | 9.85% | 10.13% |
| Day 2 | 12.67% | 12.31% | 10.13% |
| Day 3 | 11.33% | 12.92% | 12.27% |
| Day 4 | 14% | 13.54% | 11.73% |
| Day 5 | 12.67% | 9.23% | 12.27% |
| Mean | 12% | 11.57% | 11.31% |

**Note**: The body weight of fishes in each pond is listed below the pond name. The shrimp ingestion rate of fish is quantified by the daily feeding rate (weight of shrimps ingested by fish per day / fish body weight). The mean weight of dead shrimps used in the experiment is 3.9 g.

**Table S9.**

Daily healthy shrimp ingestion of grass carp (*Ctenopharyngodon idellus*) of body weight around 1 kg

| | Pond A | Pond B | Pond C | Pond D (Negative Control) |
|---|---|---|---|---|
| Fish weight | 1.05 kg | 0.956 kg | 1.013 kg | n/a |
| Number of healthy shrimps | 750 | 750 | 750 | 750 |
| Number of healthy shrimps reduced | 63 | 61 | 67 | 24 |
| Number of healthy shrimps ingested | 39 | 37 | 43 | n/a |
| Number of healthy shrimps ingested (daily) | 3.9 | 3.7 | 4.3 | n/a |
| Body weight of healthy shrimps ingested (daily) | 20.67g | 19.61g | 22.79g | n/a |
| Daily ingestion rate of healthy shrimps | 1.97% | 2.05% | 2.25% | n/a |

**Note:** In total, 750 WSSV-free shrimps (average 5.3 g of body weight) were delivered into each concrete pond. Grass carps with body weight of 1.05 kg, 0.956 kg, and 1.013 kg were put in each experiment pond separately. No fish was put in the control pond. Live shrimps remained in each pond was counted and weighted after 10 days of the experiment. The shrimp ingestion rate of fish is quantified by the daily ingestion rate (weight of shrimps ingested by fish per day / fish body weight). The number of healthy shrimps ingested by the fish is calculated by subtracting the number of healthy shrimps reduced in Pond D (negative control) from the number of healthy shrimps reduced in each of the experimental pond.

**Table S10.**

Daily healthy shrimp ingestion of African sharptooth catfish (*Clarias gariepinus*)

| | Pond A | Pond B (Negative Control) |
|---|---|---|
| Fish weight | 1050g | n/a |
| Number of healthy shrimps | 300 | 300 |
| Number of healthy shrimps reduced | 45 | 21 |
| Number of healthy shrimps ingested | 24 | n/a |
| Number of healthy shrimps ingested (daily) | 4.8 | n/a |
| Body weight of healthy shrimps ingested (daily) | 10.56g | n/a |
| Daily ingestion rate of healthy shrimps | 1.01% | n/a |

**Note:** In total, 750 WSSV-free shrimps (average 2.2 g of body weight) were delivered into each concrete pond. Grass carps with body weight of 1.05 kg was put in the experiment pond. No fish was put in the control pond. Live shrimps remained in each pond was counted and weighted after 10 days of the experiment. The shrimp ingestion rate of fish is quantified by the daily ingestion rate (weight of shrimps ingested by fish per day / fish body weight). The number of healthy shrimps ingested by the fish is calculated by subtracting the number of healthy shrimps reduced in Pond D (negative control) from the number of healthy shrimps reduced in each of the experimental pond.

**Table S11.**

Daily healthy shrimp ingestion of red drum of body weights around 0.5 kg

| | Pond A | Pond B | Pond C | Pond D (Negative Control) |
|---|---|---|---|---|
| Fish weight | 0.519 kg | 0.554 kg | 0.595 kg | n/a |
| Number of healthy shrimps | 750 | 750 | 750 | 750 |
| Number of healthy shrimps remained | 618 | 648 | 596 | 702 |
| Number of healthy shrimps reduced | 132 | 102 | 154 | 48 |
| Number of healthy shrimps ingested | 84 | 54 | 106 | n/a |
| Number of healthy shrimps ingested (daily) | 12 | 7.7 | 15.1 | n/a |
| Body weight of healthy shrimps ingested (daily) | 32.4g | 20.79g | 40.77g | n/a |
| Daily ingestion rate of healthy shrimps | 6.24% | 4.01% | 7.86% | n/a |

**Note:** In total, 750 WSSV-free shrimps (average 2.7 g of body weight) were delivered into each concrete pond. Grass carps with body weight of 0.519 kg, 0.554 kg, and 0.595 kg were put in each experiment pond separately. No fish was put in the control pond. Live shrimps remained in each pond was counted and weighted after 10 days of the experiment. The shrimp ingestion rate of fish is quantified by the daily ingestion rate (weight of shrimps ingested by fish per day / fish body weight). The number of healthy shrimps ingested by the fish is calculated by subtracting the number of healthy shrimps reduced in Pond D (negative control) from the number of healthy shrimps reduced in each of the experimental pond.

**Table S12.**

Feeding selectivity of healthy, infected (endopod and exopod removed) and dead shrimps of grass carp

| | Dead Shrimps | Infected (endopod and exopod removed) Shrimps | Healthy shrimps |
|---|---|---|---|
| Day 1 | 5.1% | 1.1% | 0. 9% |
| Day 2 | 4.9% | 1.3% | 0.2% |
| Day 3 | 5.1% | 1.1% | 0.7% |
| Day 4 | 4.7% | 1.1% | 0.7% |
| Day 5 | 4.9% | 1.3% | 0.9% |
| Day 6 | 5.1% | 2% | 0 |
| Day 7 | 5.3% | 1.6% | 0.2% |
| Day 8 | 5.5% | 1.6% | 0.4% |
| Day 9 | 4.9% | 1.3% | 0.4% |
| Mean | 5.1% | 1.4% | 0.5% |

**Note:** The WSSV-infected shrimps would die within one day. In addition, the activity of shrimps reduced after the endopod and exopod removed, which are similar to the reduction of activity of WSSV-infected shrimps. Thus, the endopod and exopod removed shrimps were used to resemble WSSV-infected shrimps. The shrimp ingestion rate of fish is quantified by the daily ingestion rate (weight of shrimps ingested by fish per day / fish body weight). The body weight of grass carp is 1.58 kg. The mean weight of shrimps used in the experiment is 3.5 g.

**Table S13.**

Feeding selectivity of healthy, infected (endopod and exopod removed) and dead shrimps of African sharptooth catfish

| | Dead Shrimps | Infected (endopod and exopod removed) Shrimps | Healthy shrimps |
|---|---|---|---|
| Day 1 | 2.4% | 0.8% | 1.6% |
| Day 2 | 2.4% | 0.8% | 0.8% |
| Day 3 | 3.2% | 1.6% | 0% |
| Day 4 | 2.4% | 0.8% | 0% |
| Day 5 | 2.4% | 0.8% | 0.8% |
| Day 6 | 1.6% | 0.8% | 0.8% |
| Day 7 | 2.4% | 0 | 0 |
| Day 8 | 2.4% | 0.8% | 0 |
| Day 9 | 1.6% | 0 | 0.8% |
| Mean | 2.31% | 0.71% | 0.53% |

**Note:** The WSSV-infected shrimps would die within one day. In addition, the activity of shrimps reduced after the endopod and exopod removed, which are similar to the reduction of activity of WSSV-infected shrimps. Thus, the endopod and exopod removed shrimps were used to resemble WSSV-infected shrimps. The shrimp ingestion rate of fish is quantified by the daily ingestion rate (weight of shrimps ingested by fish per day / fish body weight). The body weight of African sharptooth catfish is 1.03 kg. The mean weight of shrimps used in the experiment is 8.4 g.

**Table S14.**

Standard body weights of grass carps that are capable of controlling WSS outbreak determined by experiments

| Body weight (kg) | Number of shrimps | Number of live shrimps in 13 days of post-infection | Mean of survived shrimps (piece) | Standard deviation | Rate of survival (%) |
|---|---|---|---|---|---|
| 0.3 | 600 | 0 | 0 | 0 | 0 |
| | 600 | 0 | | | 0 |
| | 600 | 0 | | | 0 |
| 0.5 | 600 | 0 | 0 | 0 | 0 |
| | 600 | 0 | | | 0 |
| | 600 | 0 | | | 0 |
| 1 | 600 | 493 | 493 | 8 | 82 |
| | 600 | 486 | | | 81 |
| | 600 | 501 | | | 84 |
| 1.5 | 600 | 460 | 460 | 7 | 77 |
| | 600 | 467 | | | 78 |
| | 600 | 453 | | | 76 |
| Negative control | 600 | 502 | 502 | 9 | 84 |
| | 600 | 511 | | | 85 |
| | 600 | 493 | | | 82 |
| Positive control | 600 | 0 | 0 | 0 | 0 |
| | 600 | 0 | | | 0 |
| | 600 | 0 | | | 0 |

**Table S15.**

Body weights of African sharptooth catfish that are capable of controlling WSS outbreak determined by experiments

| Body weight （kg） | Initial Number of shrimps | Number of live shrimps in 14 days of post-infection | Rate of survival （%） | Number of dead shrimps remained in the pond | Rate (%) |
|---|---|---|---|---|---|
| 0.25 | 600 | 114 | 19 | 66 | 11 |
| 0.50 | 600 | 245 | 40.83 | 1 | 0.17 |
| 0.75 | 600 | 292 | 48.67 | 0 | 0 |
| 1.50 | 600 | 333 | 55.5 | 2 | 0.33 |
| Negative control | 600 | 74 | 12.33 | 138 | 23 |
| Positive control | 600 | 55 | 9.17 | 46 | 7.67 |

**Table S16.**

Mathematical modeled threshold parameters of using fish to control WSS spreads and experimental verification

| Body weight of shrimp (gram) | Number of expt. | Body weight of fish (kg) | Number of healthy shrimps | Success* (No. of infected shrimps) | Failure** (No. of infected shrimps) | Threshold of simulation*** |
|---|---|---|---|---|---|---|
| 1.3 | 1 | 1-kg | 750 | 3 | | 182 |
| | 2 | 1-kg | 750 | 6 | | |
| | 3 | 1-kg | 750 | 9 | | |
| | 4 | 1-kg | 750 | 12 | | |
| | 5 | 1-kg | 750 | 15 | | |
| | 6 | 1-kg | 750 | 18 | | |
| | 7 | 1-kg | 750 | 21 | | |
| 2.5 | 1 | 1-kg | 750 | 10 | | 88 |
| | 2 | 1-kg | 750 | 20 | | |
| | 3 | 1-kg | 750 | 30 | | |
| | 4 | 1-kg | 750 | 40 | | |
| | 5 | 1-kg | 750 | 50 | | |
| | 6 | 1-kg | 750 | 60 | | |
| | 7 | 1-kg | 750 | 70 | | |
| 5.0 | 1 | 1-kg | 750 | 50 | | 53 |
| | 2 | 1-kg | 750 | | 70 | |
| | 3 | 1-kg | 750 | | 90 | |
| | 4 | 1-kg | 750 | | 110 | |
| | 5 | 1-kg | 750 | | 120 | |
| | 6 | 1-kg | 750 | | 130 | |
| | 7 | 1-kg | 750 | | 140 | |
| 7.8 | 1 | 1-kg | 750 | 30 | | 32 |
| | 2 | 1-kg | 750 | | 40 | |
| | 3 | 1-kg | 750 | | 50 | |
| | 4 | 1-kg | 750 | | 60 | |

Note:

* the initial number of infected shrimps that can be successfully controlled by one 1-kg grass carp.

** the initial number of infected shrimps that fail to be controlled by one 1-kg grass carp.

*** the simulated max initial number of infected shrimps that can be successfully controlled by one 1-kg grass carp calculated by model 3

**Table S17.**

Numbers of grass carps of average body weight of 1kg for control of WSSV spreads in shrimp production

| Grass Carps (piece/hm$^2$) | Average Shrimp Yield (kg/hm$^2$) | S.D. | WSS-occurred-pond/Total Pond | WSS occurrence rate（%） |
|---|---|---|---|---|
| 0 | 3218 | 1171 | 4/5 | 80 |
| 45 | 3199 | 1211 | 4/5 | 80 |
| 150 | 4157 | 1188 | 2/5 | 40 |
| 225 | 3651 | 1195 | 2/5 | 40 |
| 300 | 8977 | 642 | 0/5 | 0 |
| 450 | 9391 | 525 | 0/5 | 0 |
| 600 | 8082 | 508 | 0/5 | 0 |
| 750 | 8002 | 671 | 0/5 | 0 |

Note: Fourty farm-size ponds (Area of 0.34 ± 0.04 hm$^2$/pond) were used to determine the number of grass carps required for the prevention of shrimp WSS in shrimp farming practice. These 40 ponds were divided into eight groups; each group consisted of 5 ponds. The amount of 675,000-shrimp/hm$^2$ of Pacific white shrimp (body length 0.8~ 1.0 cm) was released into ponds.

**Table S18.**

Effectiveness of the control of WSSV by grass carp in shrimp production at the demonstration farm in Maoming, Guangdong province, China

| Zone-Year | Area ($hm^2$) | Total Pond | Success Pond | | Yield ($kg/hm^2$) | Success (%) | |
|---|---|---|---|---|---|---|---|
| | | | Area ($hm^2$) | Pond | Mean ± S.D. | Area Ratio | Pond Ratio |
| A-2011 | 6.03 | 18 | 5.70 | 17 | 7332 ± 2059 | 94.53 | 94 |
| B-2011 | 11.03 | 28 | 2.73 | 8 | 1844 ± 2304 | 24.75 | 29 |
| A-2012 | 6.03 | 18 | 1.50 | 6 | 1953 ± 2188 | 24.88 | 33 |
| B-2012 | 11.03 | 28 | 11.03 | 28 | 8587 ± 1655 | 100.00 | 100 |

**Table S19.**

Effectiveness of the control of WSSV by African sharptooth catfish in shrimp production at the demonstration farm in Qinzhou, Guangxi province, China

| Zone-Year | Area ($hm^2$) | Total Pond | Success | | Yield ($kg/hm^2$) | Success (%) | |
|---|---|---|---|---|---|---|---|
| | | | Area ($hm^2$) | Pond | Mean ± S.D. | Area Ratio | Pond Ratio |
| A-2011 | 21.20 | 38 | 21.20 | 38 | 8730 ± 1187 | 100.00 | 100 |
| B-2011 | 67.00 | 57 | 5.33 | 4 | 450 ± 1420 | 7.96 | 7 |
| A-2012 | 21.20 | 38 | 21.20 | 38 | 9628 ± 1471 | 100.00 | 100 |
| B1-2012 | 27.00 | 25 | 27.00 | 25 | 6375 ± 1000 | 100.00 | 100 |
| B2-2012 | 40.00 | 32 | 4.36 | 3 | 500 ± 1900 | 10.90 | 9 |

**Table S20.**

The estimation of $a$, $b$, $c_1$ and $c_2$ under different weight $w$

| $w$ | $a$ | $b$ | $c_1$ | $c_2$ |
|---|---|---|---|---|
| 2.0g | 4.7321 | 0.5226 | 0.9798 | 1.1052 |
| 6.1g | 5.2799 | 0.4666 | 1.0753 | 1.1569 |
| 8.1g | 4.4270 | 0.4742 | 0.8950 | 1.1849 |

**Data S1.**

Shrimp production with or without co-cultured fishes at the farm in Maoming (Farm 1) from 2013 to 2019
